# Host genetics maps to behaviour and brain structure in mice

**DOI:** 10.1101/2022.06.18.495938

**Authors:** Sarah Asbury, Jonathan K.Y. Lai, Kelly C. Rilett, Zeeshan Haqqee, Benjamin C. Darwin, Jacob Ellegood, Jason P. Lerch, Jane A. Foster

**Author notes:** Correspondence to: Jane A. Foster, @jfosterlab (Twitter, Instagram, Facebook), www.jfosterlab.com.

## Abstract

Gene-environment interactions in the postnatal period have a long-term impact on neurodevelopment. To effectively model neurodevelopment in the mouse, we incorporated several validated behavioural tests to develop a behavioural pipeline that measures translationally relevant milestones of development in mice. The behavioral phenotype of 1060 wild type and genetically-modified mice was examined in parallel with structural brain imaging. The influence of genetics, sex, and early life stress on behaviour and neuroanatomy was determined using traditional statistical and machine learning methods. The results demonstrated that neuroanatomical diversity was primarily associated with genotype whereas behavioural phenotypic diversity was observed to be more susceptible to gene-environment interactions.

---

Advances in translational neuroscience have expanded our understanding of gene-environment mechanisms that underlie neurodevelopment and behaviour. Moreover, employing clinically-relevant genetic and environmental strategies to perturb development in mice allows researchers to determine the influence and interaction of genetics, sex, and early life environmental insults on the neurodevelopmental trajectory. These relationships are biologically shared with neurodevelopment in children and can provide an understanding of the biological mechanisms that may contribute to neurodevelopmental disorders (NDD).

The current study investigated the influence of genetics, sex, and early life stress on neurodevelopmental trajectory of mouse behaviour and neuroanatomy. The current study utilized well-established developmental milestones and behavioural assays to map the trajectory of three inbred (Balb/C, C57Bl/6NCrl, FVB) and one outbred (CD1) strain of mice, as well as T cell deficient mice, through knockout of the T cell receptor (TCR) β and δ chains (*TCRβ-/-δ-/-*) and *Fmr1* knock out (*FMR1-KO*) mice. Following behavioural testing, neuroanatomical differences were assessed by post-mortem brain volumetric analysis from MRI at 4 weeks of age. The experimental design is shown in Fig. 1. To facilitate tracking of individual mice, mice were tattooed at postnatal day (P) 2. Stressors included immune challenge on P3 with lipopolysaccharide (LPS) and maternal separation overnight on P9. Milestones and behavioural tests were selected to model key milestones of development in children and particularly those domains that are impacted in NDD.

**Fig. 1.**
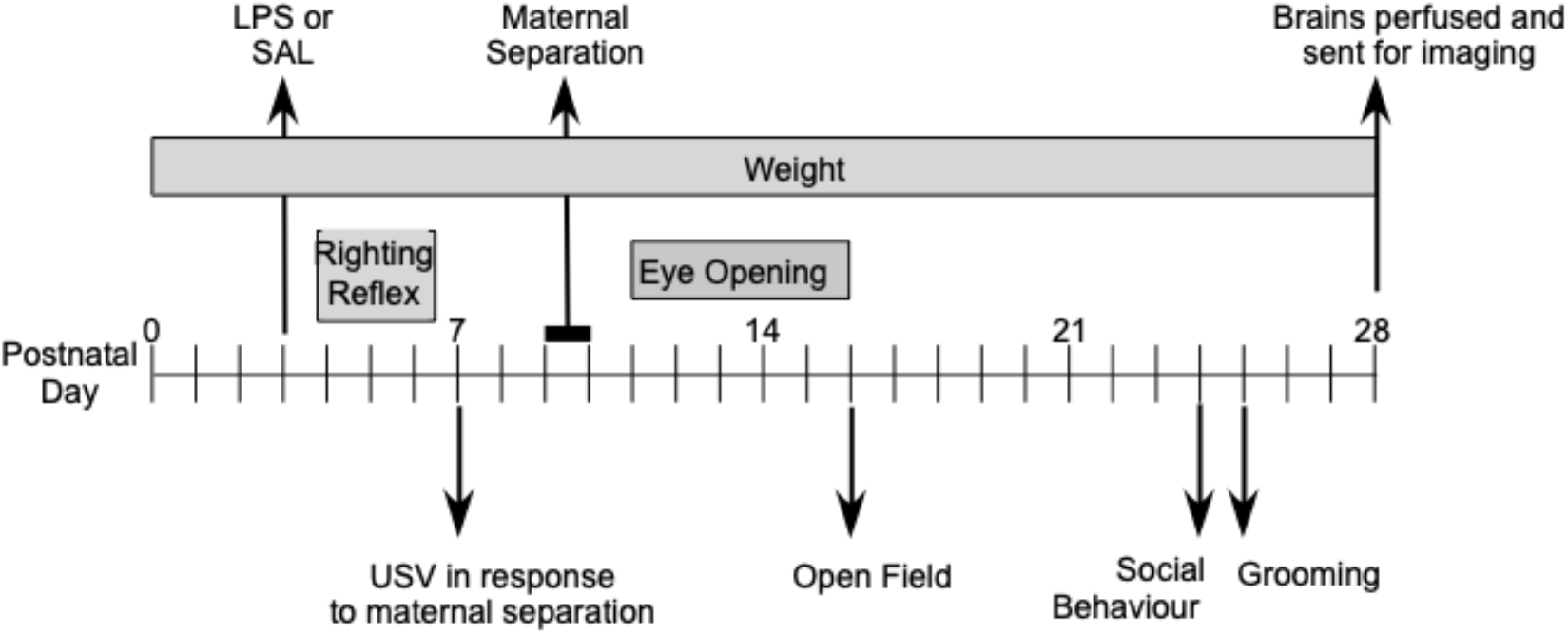
Experimental design showing postnatal challenges, developmental outcomes and behavioural tests used in the first 4 weeks of postnatal life.

Mice were assessed using our 28-day behavioural and neurodevelopmental pipeline – which included righting reflex, ultrasonic vocalizations (USV), eye opening, open field (OF), social behaviour, and self-grooming (Fig. 1). Righting reflex is a motor milestone in mice that is typically achieved between P4 to P6 (Fox, 1965; Hill et al., 2008). The time it takes for a mouse pup to flip itself on to four paws from a supine position decreases daily during this period. In response to separation from the dam, pups emit ultrasonic vocalizations. USVs peak at the end of the first postnatal week of life and provide a measure of early-life social communication (Ehret, 2005; Lai et al., 2014; Scattoni et al., 2008). Eye opening is developmental milestone that occurs between P10 and P17 (Fox, 1965; Hill et al., 2008; Scudder et al., 1967). The open field test (Roth et al.) measures exploratory behavior and locomotor activity (Brooks & Dunnett, 2009; Moy et al., 2007; Silverman et al., 2010). OF was assessed at P17 using a modified protocol that included a smaller chamber and a shorter test time to accommodate the developmental stage. Sociability was measured using the 3-chambered test (Moy et al., 2004; Moy et al., 2007; Silverman et al., 2010). Self-grooming analysis is measured to evaluate repetitive behaviour (Kalueff et al., 2016; Moy et al., 2007; Silverman et al., 2010).

The influence of genetic background on neurodevelopment and behaviour in mice has long been recognized and extensively studied across many laboratory strains using numerous behavioural and neurodevelopmental tests. Activity differences have been described between mouse strains within and beyond the strains included in our experiments (Bolivar et al., 2000; Bothe et al., 2005; Loos et al., 2014; Molenhuis et al., 2014; Tucci et al., 2006). There are also recognized differences in sociability (Moy et al., 2007), early life social communication(Lai et al., 2014), grooming (Kalueff & Tuohimaa, 2004, 2005; Molenhuis et al., 2014), and motor development (Dierssen et al., 2002; Molenhuis et al., 2014; Roth et al., 2013) between strains. In accordance with inbred mouse strain background influencing behaviour, several studies have identified candidate genes or quantitative trait loci associated with phenotypic differences for mouse neurobehavioral traits including activity (Kelly et al., 2003), rearing (Delprato et al., 2017), grooming (Delprato et al., 2017), early life social communication (Ashbrook et al., 2018) and sociability (Takahashi et al., 2010). These studies broadly demonstrate genetic determinants of normal developmental neurobehavioral phenotypes. A large-scale study of mouse behaviour and cognition demonstrated that single nucleotide polymorphisms also influence behavioural, anxiety, and cognitive outcomes in mice (Janecka et al., 2017). Such findings indicated that genetics are a major source of variability in behavioural phenotypes.

Using both traditional statistical and machine learning methods neurodevelopmental, behavioural, and neuroanatomical differences between inbred mouse strains were demonstrated with similar neurodevelopmental trajectories between knockout mice and their wild type strains. These findings validate the importance of host genetics underlying behavioural and neuroanatomical differences between mice.

## METHODS

### Animals

Breeding pairs were bred in house and pregnant dams were ordered from Charles River. Mice strains included FVB, C57Bl/6NCrl (B6), Balb/C, CD1, *FMR1-*KO (FVB.129P2.*Fmr1tm1Cgr*/J, stock #004624) and *TCRβ-/-δ-/-* on a C57Bl/6 background (Rilett et al., 2015). Mice were housed at the animal facility at St. Joseph’s Healthcare with food and water available *ad libitum,* under a 12h:12h light dark cycle with lights on at 5 AM and lights off at 5 PM. Birth was set as postnatal day 0 (P0). On P2, litters were culled to 10 pups and pups were uniquely tattooed on their paw for identification. Pups were weaned on P21 and caged by sex with up to 4 littermates per cage. All experimental procedures were approved by the Animal Research Ethics Board, McMaster University in accordance with the guidelines of the Canadian Council on Animal Care.

### Experimental Design

Our experimental design included developmental milestones, righting reflex, ultrasonic vocalizations (USVs), eye opening, open field, sociability and self-grooming. This analysis was completed over the first 28 days of life with postnatal challenges occurred on P3 (immune) and P9 (stress) (Fig. 1). Experimental mice (Table 1) included 1060 pups from 159 litters (29 FVB, 30 B6, 38 Balb/C, 28 CD1, 14 *FMR1*-KO, 20 *TCRβ-/-δ-/-*).

**Table 1:** Experimental mice used in behavioural analyses

| Strain | Sex | P3 Treatment | P9 Treatment | N |
| --- | --- | --- | --- | --- |
| **B6** | F | SAL | CON | 23 |
|  |  |  | MS | 25 |
|  |  | LPS | CON | 24 |
|  |  |  | MS | 22 |
|  | M | SAL | CON | 25 |
|  |  |  | MS | 23 |
|  |  | LPS | CON | 24 |
|  |  |  | MS | 23 |
| **TCR** | F | SAL | CON | 13 |
|  |  |  | MS | 16 |
|  |  | LPS | CON | 13 |
|  |  |  | MS | 14 |
|  | M | SAL | CON | 9 |
|  |  |  | MS | 15 |
|  |  | LPS | CON | 11 |
|  |  |  | MS | 18 |
| **BalbC** | F | SAL | CON | 25 |
|  |  |  | MS | 27 |
|  |  | LPS | CON | 32 |
|  |  |  | MS | 34 |
|  | M | SAL | CON | 21 |
|  |  |  | MS | 23 |
|  |  | LPS | CON | 23 |
|  |  |  | MS | 24 |
| **CD1** | F | SAL | CON | 31 |
|  |  |  | MS | 32 |
|  |  | LPS | CON | 29 |
|  |  |  | MS | 28 |
|  | M | SAL | CON | 36 |
|  |  |  | MS | 28 |
|  |  | LPS | CON | 36 |
|  |  |  | MS | 30 |
| **FVB** | F | SAL | CON | 28 |
|  |  |  | MS | 23 |
|  |  | LPS | CON | 24 |
|  |  |  | MS | 44 |
|  | M | SAL | CON | 21 |
|  |  |  | MS | 24 |
|  |  | LPS | CON | 24 |
|  |  |  | MS | 25 |
| **FMR1** | F | SAL | CON | 10 |
|  |  |  | MS | 10 |
|  |  | LPS | CON | 11 |
|  |  |  | MS | 11 |
|  | M | SAL | CON | 14 |
|  |  |  | MS | 11 |
|  |  | LPS | CON | 10 |
|  |  |  | MS | 13 |

### Postnatal challenges

On P3, pups were administered lipopolysaccharide (LPS) (0.1 mg/kg; *Escherichia coli* LPS; Sigma, St. Louis, MO) or saline i.p. at 50 μl/g. Injections were done at 4 PM. On P9, maternal separation (MS) or control (Loos et al.) treatment was administered. For MS litters, pups were weighed at 4 PM, and then the dam was removed from the home cage at 5 PM. The home cage was then placed on a heating pad at 37°C until 9 AM the next day, when the pups were weighed and the dam returned to the cage. Pups were again weighed at 4 PM. For control litters, pups were weighed at 4 PM on P9, 9 AM on P10 and 4 PM on P10; dams were removed briefly during weights and then returned to the home cage.

### Righting Reflex

On P4-P6, pups were tested for motor development by timing their ability to right themselves after being placed on their backs. Testing was done at 4 PM. A completed righting was defined by all four paws on the ground simultaneously. Time was kept with a stopwatch. The maximum score was set at 30 s at which time the pup was manually righted. The righting reflex score was calculated by taking the average righting reflex time from P4-P6. Lower righting reflex score therefore indicates earlier achievement of the righting reflex.

### Eye Opening

From P10 to P17, eye opening was scored daily: a score of 0, 1, or 2 was assigned per mouse reflecting the number of eyes open. Eye opening score was calculated by averaging the eye opening data for each mouse from P10 to P17. A higher eye opening score reflects earlier eye opening of one or both eyes.

### USV Recordings

On P7, pups were consecutively maternally separated from the dam and littermates and placed in a custom-made sound-attenuating chamber. Testing took place during the first half of the active period, at least one hour after the active cycle began. Ultrasonic vocalizations were recorded for 3 min and then each pup was transferred to a separate holding cage. After all pups were tested, the pups were returned to the dam. Vocalizations were digitized using an Avisoft UltraSoundGate 116-200 recording device and USG CM116/CMPA microphone. The microphone was clamped to a retort stand and situated 17.5 cm above the center of the recording chamber. Calls were digitized in real-time and subsequently analyzed with Avisoft SAS Lab Pro.

### Open Field

At P17, pups were tested in the open field. Behavioural testing was conducted in a non-colony room after a 30 min habituation to the room. Testing took place in low light during the first half of the active period. Behaviours were automatically recorded for 15 min using the Kinder Scientific Smart Rack System consisting of a 24 cm wide x 45 cm long x 24 cm high cage rack system, with 22 infrared beams (7 X & 15 Y) and a rearing option (22 additional beams). A Plexiglas^®^ box was placed at one end of the chamber to reduce the testing chamber size to 24 x 23 cm. Data were collected using MotorMonitor^®^ software (Kinder Scientific, Poway, CA). A maximum of 6 pups were tested at a time. After all pups had undergone testing, they were returned to the dam; maternal separation did not exceed 30 min.

### Sociability

At P24, sociability was tested using a 3-chamber apparatus (Moy et al., 2004; Nadler et al., 2004). Behavioural testing was conducted in a non-colony room after a 20 min habituation to the room. Testing took place during the first half of the active period. Mice were placed in the centre zone of the chamber with no access to the other chambers for 5 minutes. Subsequently, an age-, strain-, and sex-matched stranger mouse was placed in an inverted cup in one of the side chambers, the doors from the centre chamber to the outer chambers opened, and behaviour was recorded for 10 minutes. Live-tracking and automated videotape analysis was done using EthoVision® software.

### Self-grooming

At P25, mice were observed for 10 min in a standard housing cage without bedding and scored for time spent grooming (Silverman et al., 2010). Mice were habituated to the testing cage for 10 min prior to grooming test. Grooming behaviours were manually scored using AnyMaze software. Grooming data was only included if inter-rater reliability was demonstrated (κ >0.05).

### Perfusions

Mice were perfused at P28 at St. Joseph’s Healthcare in Hamilton, Ontario prior to being transferred to the Mouse Imaging Centre in Toronto for imaging and analysis. The perfusion protocol was as follows: Mice were anesthetized with ketamine/xylazine and intracardially perfused with 30 mL of 0.1 M PBS containing 10 U/mL heparin and 2 mM ProHance (Bracco Diagnostics Ltd. a Gadolinium contrast agent) followed by 30 mL of 4% paraformaldehyde (PFA) containing 2 mM ProHance (Cahill et al., 2012). Perfusions were performed with a minipump at a rate of approximately 1mL/min. After perfusion, mice were decapitated and the skin, lower jaw, ears, and the cartilaginous nose tip were removed. The brain and remaining skull structures were incubated in 4% PFA + 2 mM ProHance overnight at 4°C then transferred to 0.1M PBS for at least 1 month prior to MRI scanning (de Guzman et al., 2016).

### Behavioural analysis

Analysis of behavioural data to identify main effects of genotype, sex, and age was completed in SPSS Ver. 25 by multivariate general linear models followed by Bonferroni-corrected posthoc tests. Omnibus analysis was conducted in behavioural data prior to P9 (RR, USV) that included 2 treatment (P3) groups (LPS, SAL) and in behavioural data following P9 (eye opening, OF, sociability, self-grooming) that included 4 treatment (P3_P9) groups (SALCON, SALMS, LPSCON, LPSMS). Based on main effects and interactions identified in the omnibus analyses, behaviour test-specific multivariate (genotype, sex, treatment – all test outcomes) and univariate analysis (genotype, sex –individual test outcomes) was conducted (SPSS Ver. 25) followed by pairwise comparisons using Bonferonni-corrected posthoc tests.

### MRI Analysis

All images were acquired using a 7.0 Tesla MRI Scanner (Agilent Inc., Palo Alto, CA). For neuroanatomical scans, a 40 cm inner bore diameter gradient was used with a maximum gradient strength of 30 G/cm. This was used in conjunction with a custom built solenoid array capable of scanning 16 brains at a time (Dazai et al., 2011; Lerch et al., 2011). To assess the volume differences throughout the brain, a T2-weighted 3D fast spin echo (FSE) sequence is used that is optimized for gray/white matter contrast. Parameters for the sequence: TR of 2000 ms, and TEs of 10 ms per echo for 6 echoes, with the centre of k-space being acquired on the 4^th^ echo, TE_eff_ of 42ms, two averages, field-of-view (FOV) of 14 × 28 × 25 mm and matrix size of 250 × 504 × 450 giving an image with 0.056mm isotropic voxels. In the first phase encode direction consecutive lines of k-space were assigned to alternating echoes to move discontinuity related ghosting to the edges of the FOV (Thomas et al., 2004). This sequence involves oversampling k-space in the first phase encode direction by a factor of 2 to avoid ghosting in the final image. This gives a FOV of 28 mm but is subsequently cropped to 14 mm after reconstruction. Total imaging time was approximately 12 hours (Lerch et al., 2011).

To visualize and compare differences between mouse brains the images are registered together (both linear and nonlinear registration). All scans were then resampled with the appropriate transform and averaged to create a population atlas representing the average anatomy of the study sample. All registrations were performed with a combination of mni_autoreg tools (Collins et al., 1994) and advanced normalization tools (ANTs) (Avants et al., 2008; Avants et al., 2011). The result of the registration was to have all scans deformed into alignment with each other in an unbiased fashion. This allowed for the analysis of the deformations needed to take each individual mouse’s anatomy into this final atlas space. Allowing modelling of the deformation fields as they relate to genotype (Lerch et al., 2008; Nieman et al., 2007). Significant volume changes were calculated by warping a pre-existing classified MRI atlas onto the population atlas, which allowed for the volume of 159 segmented structures encompassing cortical lobes, large white matter structures (i.e. corpus callosum), ventricles, cerebellum, brain stem, and olfactory bulbs to be assessed in all brains. This classified atlas incorporated the structures from three separate pre-existing atlases: 1) which delineated 62 different regions throughout the brain (Dorr et al., 2008), 2) which further classified multiple different areas in the cerebellum (Steadman et al., 2013), and 3) which divided the cortex into 64 different regions (Ullmann et al., 2013). Moreover, these measurements were examined on a voxel-wise basis in order to localize the differences found within regions or across the brain. Regional and voxelwise differences were examined using two measures, absolute volume (mm^3^) as well as relative volume (% total brain volume). Multiple comparisons in this study were controlled for using the False Discovery Rate (Genovese et al., 2002).

Brain regions – as defined using the Dorr-Steadman-Ullmann-Richards-Qiu-Egan (40 micron, DSURQE) atlas and Allen Brain Institute hierarchy – were used for PCA, clustering and Random Forest analyses (Lerch et al., 2017). Hierarchical brain regions were pruned to level 8 for all regions for most regions. Hippocampus was instead pruned to level 7 and Midbrain was pruned to level 9. Only regions represented by terminal nodes were used for downstream analyses. Regions with absolute volume >1 mm^3^ were removed. The regions used in downstream analyses are available as a list in Supplemental File 1 or as a hierarchical visualization (Supplementary Figure S1).

## INTEGRATIVE ANALYSIS OF BEHAVIOURAL DATA

### Principle coordinate analysis

PCA and hierarchical clustering analyses were performed using *prcomp*, *hclust,* and *dist* in the stats R package (R Core Team, 2021). Z-score normalized behavioural outcomes: average time for righting reflex (P4-P6); USV calls, duration, and intercall interval; average eye opening score; open field rearing; time spent in mouse, center, and empty chamber in three chamber test; and grooming frequency, duration, and latency were analyzed for behavioural PCA. Z-score normalized MRI relative volumes from postnatal day 28 were used for neuroanatomical PCA. Brain regions used for analysis are listed in Supplementary file 3. Hierarchical clustering was performed on all neurodevelopmental and behavioural or MRI data using Euclidian distances and Ward.D2 clustering method. 3D PCA visualizations were generated using RGL R package (Adler, 2020) and *scatter3D* from car R package (Fox, 2019). For behavioural networks analysis, the optimal number of clusters (k) was determined using WSS and silhouette plots generated using Factoextra R package (Kassambara, 2020). Clustering results were then visualized using the dendrogram function in base R, and separated into k = 2 groups using cutree function in base R. Cluster stability for behavioural and neurodevelopmental, neuroanatomical, or behavioural network clustering was determined using fpc package *clusterboot* (Henning, 2020). Jaccard mean > 0.75 is considered a stable cluster, 0.6 – 0.75 is considered a pattern in the data, 0.5 – 0.6 is considered an unstable cluster and < 0.5 is considered a dissolved cluster.

### Random Forest Models

Random Forest models predicting genotype, sex, or combined treatment from behaviour and neurodevelopmental outcomes were generated using the following predictor variables: average time for righting reflex (P4-P6), USV calls, USV duration, USV intercall interval, average eye opening score, OF rearing, OF total distance, 3-chambered sociability center chamber time, 3-chambered sociability mouse chamber time, 3-chambered sociability empty chamber time, grooming frequency, grooming latency, grooming duration. Sex, P3 and P9 treatment, or genotype was also included as predictor variables for the respective models. Models predicting genotype using relative neuroanatomical volume used predictor variables listed in Supplementary File 1. Random Forest models were generated using the randomForest package in R (Liaw, 2002). Ten test and validation sets were generated via random 80/20 splits. Test sets were further segmented via random 80/20 split for 5-fold cross-validation hyper parameter tuning of the number of trees (500 – 5000) and number of predictor variables available at each split (behaviour predictor variables mtry = 1:7 or neuroanatomical predictor variables mtry = 1:10). Model accuracy was reported as validation set results using Misclassification Error (ME) or Adjusted Rand Index (ARI) calculated using the e1071 package (Meyer, 2020). Predictor variable importance was measured via mean decreased Gini index for the best performing set. When the random forest model accurately predicts the response variable, high predictor variable importance rank implies an association between the predictor variable and response variable.

## RESULTS

### Behavioural trajectories and developmental milestones

Representative graphs of the behavioural results are provided for male and female C57Bl/6 mice in Fig. 2 and Fig. 3. Righting reflex develops between P4 and P6 as shown in Fig. 2A and the average time to right is also provided. Refer to supplemental file 2 – descriptive statistics for P3 and P3_P9 analyses.

**Fig. 2.**
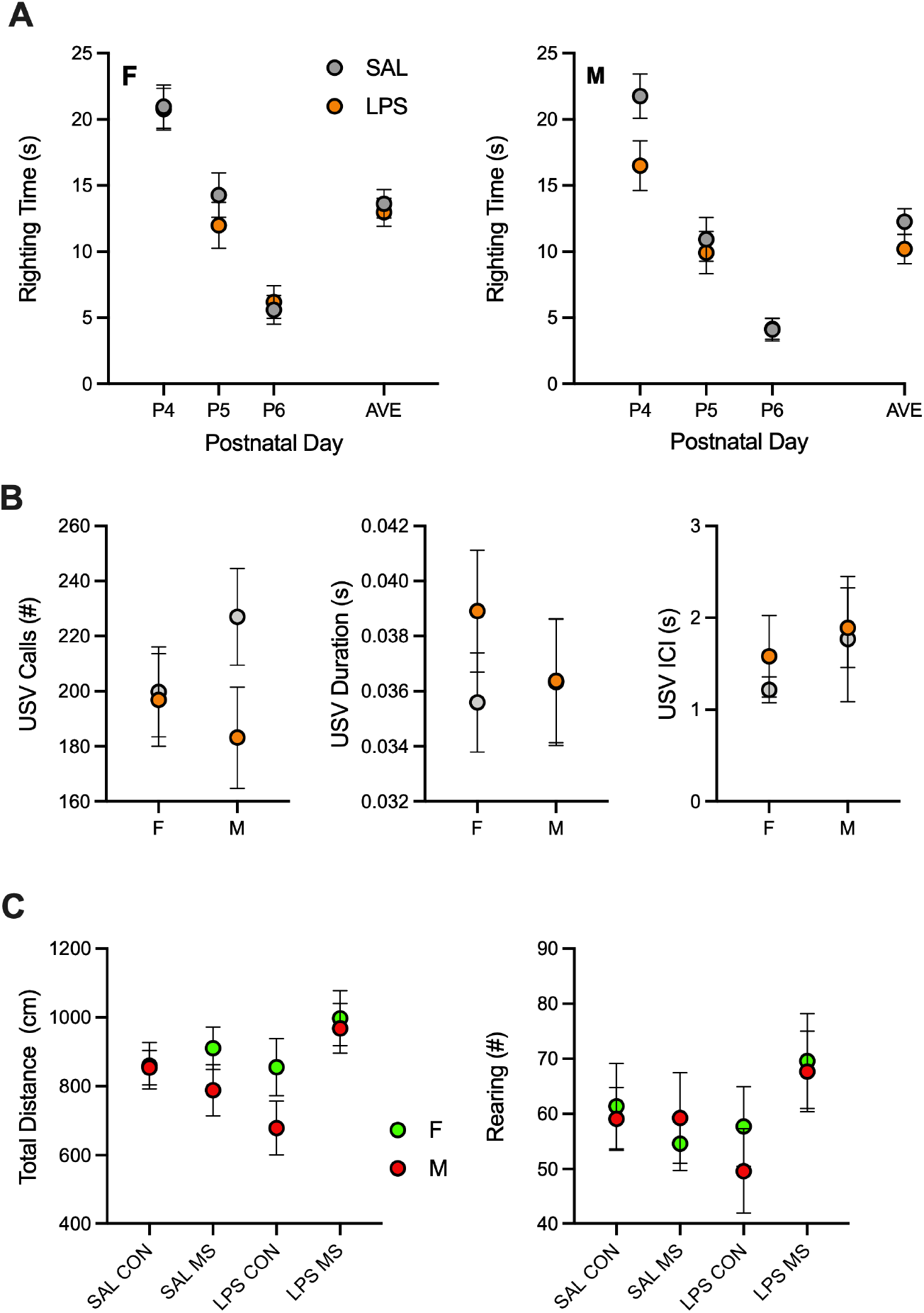
Representative behavioural graphs for righting reflex on postnatal day (P) 4-6 and average righting reflex time (A), ultrasonic vocalizations (USV) (B), SAL treated mice are shown as grey and LPS treated mice as orange (A,B). Total distance travelled in the Open Field at P17 is shown for different treatment conditions at P3 and P9 - SALCON, SALMS, LPSCON, LPSMS (C). Data is shown for female and male C57Bl/6 mice (mean +/- S.E.).

**Fig. 3.**
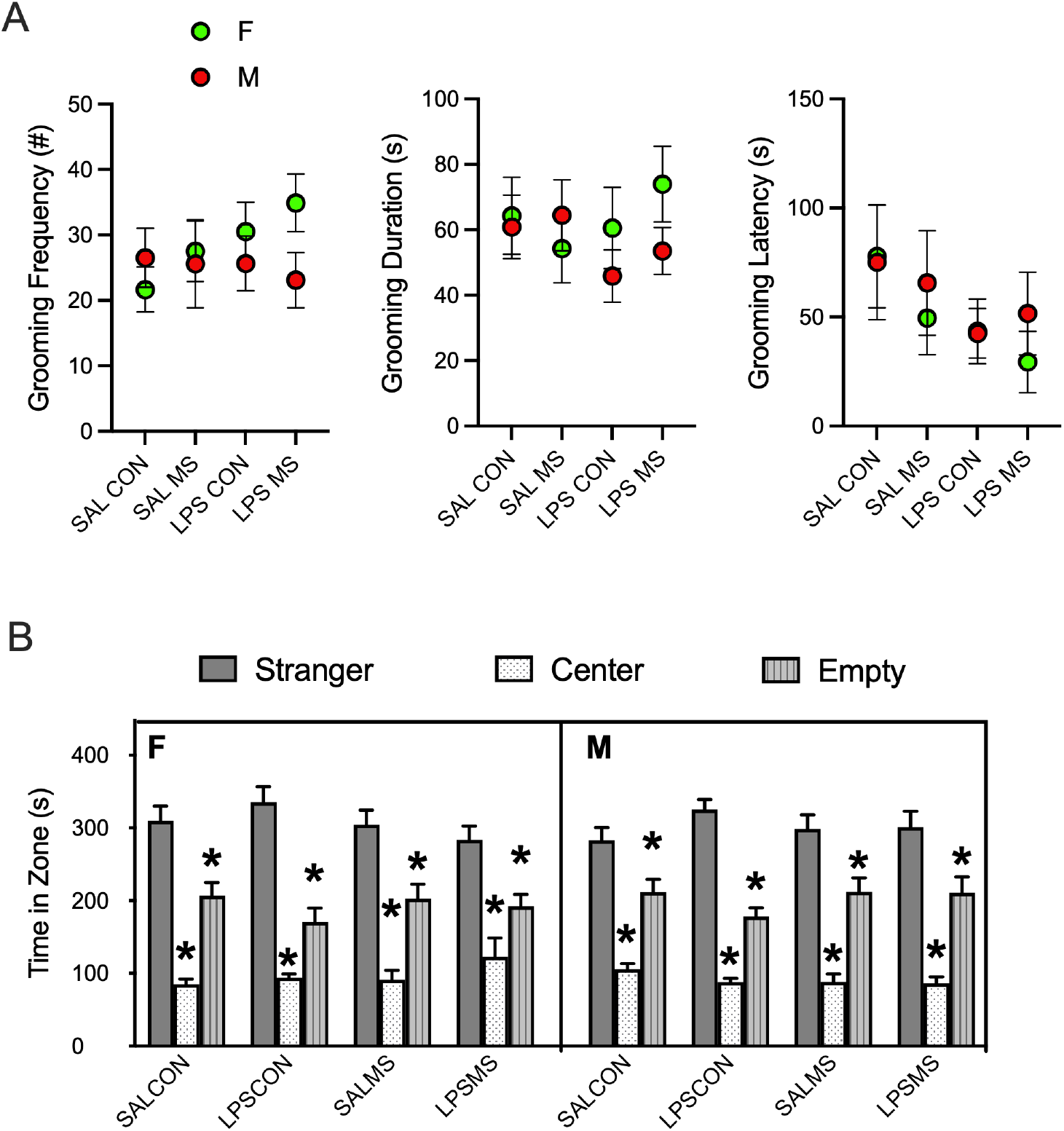
Representative behavioural graphs for self grooming (A) and sociability (B). Data is shown for female and male C57Bl/6 mice (mean +/- S.E.). Female (B-left panel) and male (B-right panel) mice showed typical sociability measured as social preference for chamber with stranger mouse. * p<0.05, significantly different from stranger time.

### ANOVA and posthoc behavioural analysis

Genotype, sex, and treatment effects and interactions were investigated using multivariate and univariate ANOVAs for each neurodevelopmental and behavioural outcome. Neurodevelopmental and behavioural differences were primarily genotype effects. There were no independent sex effects for between-subject effects ANOVA and limited independent treatment effects (Table 2).

**Table 2:** Multivariate ANOVA investigating association of genotype, sex, and treatment with <u>behaviour</u>

| Test | Parameter | Effect | df | F | Sig. |
| --- | --- | --- | --- | --- | --- |
| Righting Reflex | Average Righting Reflex | GENOTYPE | 5 | 130.2 | <0.001 |
|  |  | GENOTYPE * SEX | 5 | 3.8 | 0.002 |
| USV | USV Calls | GENOTYPE | 5 | 127.6 | <0.001 |
|  | USV Duration | GENOTYPE | 5 | 115.3 | <0.001 |
|  |  | GENOTYPE * Treatment<sup>1</sup> | 5 | 2.5 | 0.027 |
|  | USV ICI | GENOTYPE | 5 | 60.6 | <0.001 |
| Eye Opening | Eye Opening Score | GENOTYPE | 5 | 30.0 | <0.001 |
| Open Field | OF Rearing | GENOTYPE | 5 | 69.1 | <0.001 |
|  |  | Combined Treatment | 3 | 6.8 | <0.001 |
|  |  | GENOTYPE * Treatment | 15 | 2.2 | 0.005 |
|  |  | GENOTYPE * SEX | 5 | 3.8 | 0.002 |
|  | OF Total Distance | GENOTYPE | 5 | 86.5 | <0.001 |
|  |  | Combined Treatment | 3 | 4.5 | 0.004 |
|  |  | GENOTYPE * Treatment | 15 | 2.8 | <0.001 |
|  |  | GENOTYPE * SEX | 5 | 5.4 | <0.001 |
| Sociability | Center Chamber | GENOTYPE | 5 | 77.2 | <0.001 |
|  | Empty Chamber | GENOTYPE | 5 | 3.9 | 0.002 |
|  | Mouse Chamber | GENOTYPE | 5 | 31.9 | <0.001 |
|  |  | SEX * Treatment | 3 | 3.0 | 0.031 |
| Grooming | Grooming duration | GENOTYPE | 5 | 43.3 | <0.001 |
|  | Grooming frequency | GENOTYPE | 5 | 29.9 | <0.001 |
|  |  | GENOTYPE * SEX | 5 | 2.8 | 0.016 |
|  |  | SEX * Treatment | 3 | 3.5 | 0.015 |
|  | Grooming latency | GENOTYPE | 5 | 7.3 | <0.001 |
|  |  | GENOTYPE * SEX | 5 | 3.3 | 0.006 |

<sup>1</sup> Treatment only refers to P3 treatment (LPS or SAL), as developmental milestone is assessed before P9 treatment (CON or MS).

The average righting reflex neurodevelopmental metric measures the average righting reflex time between P4 and P6. Low average righting reflex values indicates faster development of the righting reflex neurodevelopmental milestone. Genotype was a main effect for average righting reflex (p < 0.001) (Table 2), and posthoc analysis indicates that the FVB strain has earlier development of the righting reflex compared to other wildtype inbred genotypes (Table 3). *TCRβ-/-δ-/-* mice also have significantly higher average righting reflex time compared to wildtype B6 (p < 0.001), indicating delayed righting reflex development in T-cell deficient mice (Table 3).

**Table 3:**
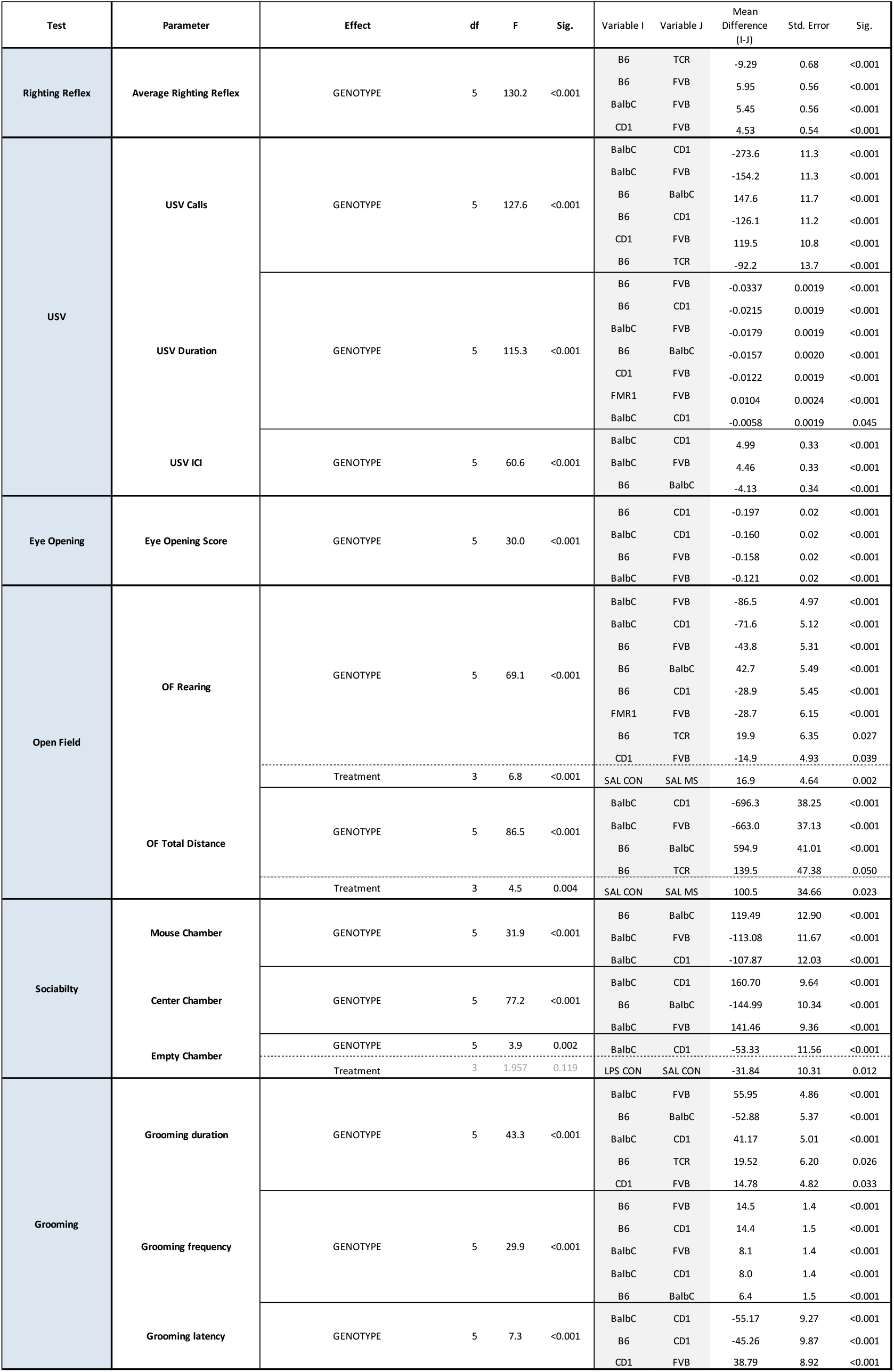
Behaviour univariate ANOVA with Bonferroni-corrected posthoc

USVs measure early life social communication. Genotype was a significant main effect for USV call number (p < 0.001) (Table 2). Most strains and knockout mice having significant pair-wise differences (Table 3). For wildtype mice, Balb/C mice have the lowest USV call number, while CD1 mice have the highest USV call number (Table 3). USV call duration was shortest in B6 mice and longest in FVB mice (Table 3). USV intercall intervals had a significant genotype effect (p < 0.001), driven by high intercall intervals of Balb/C mice compared to other wildtype strains, in accordance with lowest number of USV calls (Table 3). *TCRβ-/-δ-/-* mice have a higher number of calls compared to wildtype B6 mice (p < 0.001), however T cell knockout did not affect USV duration (Table 3). Contrastingly, *FMR1-KO* mice have a lower number of USV calls compared to wildtype FVB mice (p < 0.001), with no differences in USV duration (Table 3).

Higher eye opening score indicates the eye opening milestone occurred earlier in development. Eye opening score was dependent on genotype (p < 0.001) (Table 2) and lowest in B6 and Balb/C mice (Table 3), indicating these strains attain eye-opening milestone later in development. In contrast, CD1 and FVB have the highest eye-opening scores (Table 3). There were no knockout effects in eye opening score (Table 3).

Balb/C mice have pronouncedly lower activity metrics in the open field test; both total distance and rearing scores were lower than other strains (Table 3). Rearing frequency was highest in FVB mice, followed closely by CD1 mice (Table 3). Notably, *FMR1-KO* mice had a higher rearing frequency compared to their background strain FVB (p < 0.001) (Table 3). Furthermore, rearing and open field total distance were lower in *TCRβ-/-δ-/-* mice compared to their wildtype B6 counterparts (p = 0.027, p = 0.050), indicating decreased activity in T-cell deficient mice (Table 3). Open field total distance was similar between the wildtype genotypes, except for Balb/C mice which had markedly lower activity (Table 3). Both open field total distance and rearing had a treatment effect (p < 0.001, p = 0.005) (Table 2) driven by reduced activity in maternally separated mice that were not LPS-treated compared to controls (Table 3).

Sociability, as measured by time spent in the mouse chamber, had a significant genotype effect (p < 0.001) and was lowest in Balb/C mice but similar between wildtype genotypes (Table 3). There were no significant differences in sociability for knockout mice (Table 3). LPS treated mice that were not maternally separated spent less time in the empty chamber according to posthoc analysis, however treatment effects were not detected for empty chamber time via between-subject ANOVA (p = 0.119) indicating the detected posthoc difference was coincidental.

Finally, grooming duration was highest in Balb/C mice, although CD1 mice also had high grooming duration relative to other strains (Table 3). While grooming frequency is highest in B6 mice, Balb/C also have higher grooming frequency compared to most other strains (Table 3). Balb/C mice have a grooming phenotype of longer frequent grooms, B6 mice have a phenotype of short and highly frequent grooms. For latency to first groom, CD1 have the highest metric compared to other wildtype mice (Table 3). The only knockout effect for grooming behaviours is decreased grooming duration in *TCRβ-/-δ-/-* compared to wildtype B6 (p = 0.026) (Table 3).

### Principle coordinate cluster analysis of behaviour

Inbred mice overall behavioural and neurodevelopmental phenotypes were visualized using PCA plots (Fig. 4A). Inbred mouse strains cluster based on neurodevelopmental and behavioural outcomes. We used unsupervised hierarchical clustering to form 3 clusters. Each unsupervised cluster predominantly maps to a single inbred strain (Fig. 4B). The structure of the clusters demonstrates a robust relationship between genetic background and the overall behavioural and neurodevelopmental phenotype of mice.

**Figure 4:**
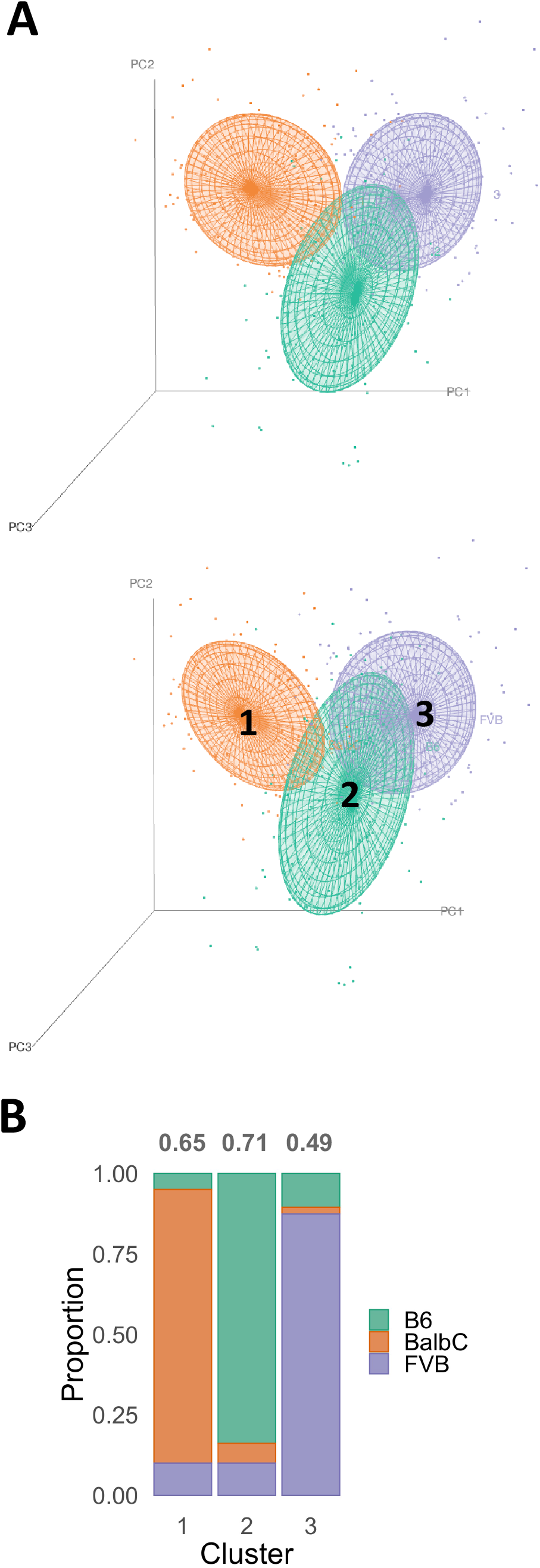
Principal component analysis of all neurodevelopment and behavioral metrics in inbred mice. **A) Top:** PCA points coloured by genotype. Balb/C = Orange, FVB = Purple, B6 = Green. **Bottom:** PCA points coloured by 3 clusters generated by unsupervised hierarchical clustering (k = 3). Orange = Cluster 1 (Jaccard Index = 0.65), Green = Cluster 2 (Jaccard Index = 0.71), Purple = Cluster 3 (Jaccard Index = 0.49). **B)** Behavioural and neurodevelopmental hierarchical cluster membership by mouse strain. Cluster stability is annotated above the bar graph for each cluster. Cluster stability was measured by the average Jaccard similarity index with the clusters of 1000 bootstrapped samples.

When *FMR1-KO* mice are included for unsupervised hierarchical clustering, all *FMR1-KO* mice were members of the behavioural and neurodevelopmental cluster associated with their background strain, FVB (Supplementary Fig. S2). *FMR1-KO* mice do not cluster with the Balb/C or B6 dominant clusters. While differences exist between FVB and *FMR1-KO* mice in specific USV and OF metrics, their overall behavioural and neurodevelopmental phenotype retains higher similarity to FVB mice than other inbred strains in our study. However, the inclusion of *FMR1-KO* mice in the PCA plot decreases the effectiveness of hierarchical clustering discernment of FVB and B6 mice when there are three unsupervised clusters. Although the percent composition of each B6 and FVB predominant clusters remains similar in both analyses, 36 B6 mice shift to the FVB cluster when *FMR1-KO* mice are included. Changes in the B6 dominant cluster were an artefact of the sensitivity of hierarchical clustering to separating the overlap between the B6 and FVB behavioural phenotypes, whereby FVB mice with a more distant phenotype from the average mouse in the FVB clusters were assigned to the B6 cluster.

Similarly, when *TCRβ-/-δ-/-* mice are included in the PCA inbred plots, their behavioural and neurodevelopmental phenotype overlaps with their background strain B6 (Supplementary Fig. S2). They cluster primarily with B6 mice, with few *TCRβ-/-δ-/-* mice assigned to the FVB predominant cluster. There are more behavioural and neurodevelopmental differences of T-cell knockout mice compared to their background strain than *FMR1-KO* and FVB mice. Differences between B6 and *TCRβ-/-δ-/-* neurodevelopmental and behavioural metrics include: righting reflex, USV, OF, and grooming. Despite this, *TCRβ-/-δ-/-* mice still predominantly cluster with their background strain, albeit with lower cluster purity compared to the *FMR1-*KO and FVB mice cluster.

The addition of the outbred heterozygous strain CD1 disrupts the hierarchical clustering such that the B6 and FVB clusters are dissolved (Supplementary Fig. S3). The clustering pattern of CD1 mice suggests their behavioural and neurodevelopmental phenotype varies on a phenotypic spectrum that intersects with B6 and FVB mice, creating overlap that dissolves the mice into a single cluster. The remaining portion of the B6 predominant cluster begins to cluster with a portion of the Balb/C predominant cluster (Supplementary Fig. S3). These clustering patterns reflect that outbred heterozygous CD1 mice have heterogenous behavioural and neurodevelopmental phenotypes that span B6 and FVB trajectories.

The influence of sex and treatment on behavioural and neurodevelopmental clustering of mice was also assessed. There was no appreciable cluster separation of any sex-treatment subgroup (Supplementary Fig. S4).

### Random Forest Genotype Prediction from Behavioural and Neurodevelopmental milestones

Accurate prediction of genotype from behavioural and neurodevelopmental milestones further validates distinct phenotypes between genotypes and inbred strains of mice in our study. Random Forest models were trained to predict mouse genotype using sex, treatment, and behavioural and neurodevelopmental outcomes. The average genotype prediction accuracy of our model was 73% across 10 training/test/validation sets (Misclassification Error (ME) = 0.27±0.02, Adjusted Rand Index (ARI) = 0.49±0.03). 73% accuracy of machine learning models for predicting genotype from behaviour reinforces that the structure of behavioural and neurodevelopmental data differs between mouse genotypes and strains – indicative of unique phenotypes. A breakdown of prediction accuracy for each genotype is reported as a proportionate classification matrix (Figure 5). As demonstrated in PCA and hierarchical clustering analysis, inbred strains can be accurately distinguished from one another for most observations (Fig. 3, Fig. 5A). However, behavioural and neurodevelopmental phenotypic overlap often occurs between the outbred CD1 mice, or knockout mice and their background strains (Fig. 5A, Fig S2). CD1 mice are most frequently misclassified as FVB or B6 mice, akin to the results when CD1 were included for hierarchical clustering (Fig. 5A). FVB misclassifications are predominantly misclassified as their associated genetic knockout – *FMR1-KO* – in 20% of cases, or as outbred CD1 strain in 14% of cases. B6 genotype misclassifications are predominantly misclassification as CD1 or *TCRβ-/-δ-/-* genotypes. In accordance with observed overlap between knockout mice and their background strain in hierarchical clustering and PCA visualization method, *TCRβ-/-δ-/-* and *FMR1-*KO misclassification is almost solely attributable to misclassification as their wildtype background strains. Balb/C mice do not demonstrate appreciable misclassification as any other genotype, reaffirming their distinct behavioural and neurodevelopmental phenotype in our study (Fig. 5A).

**Figure 5:**
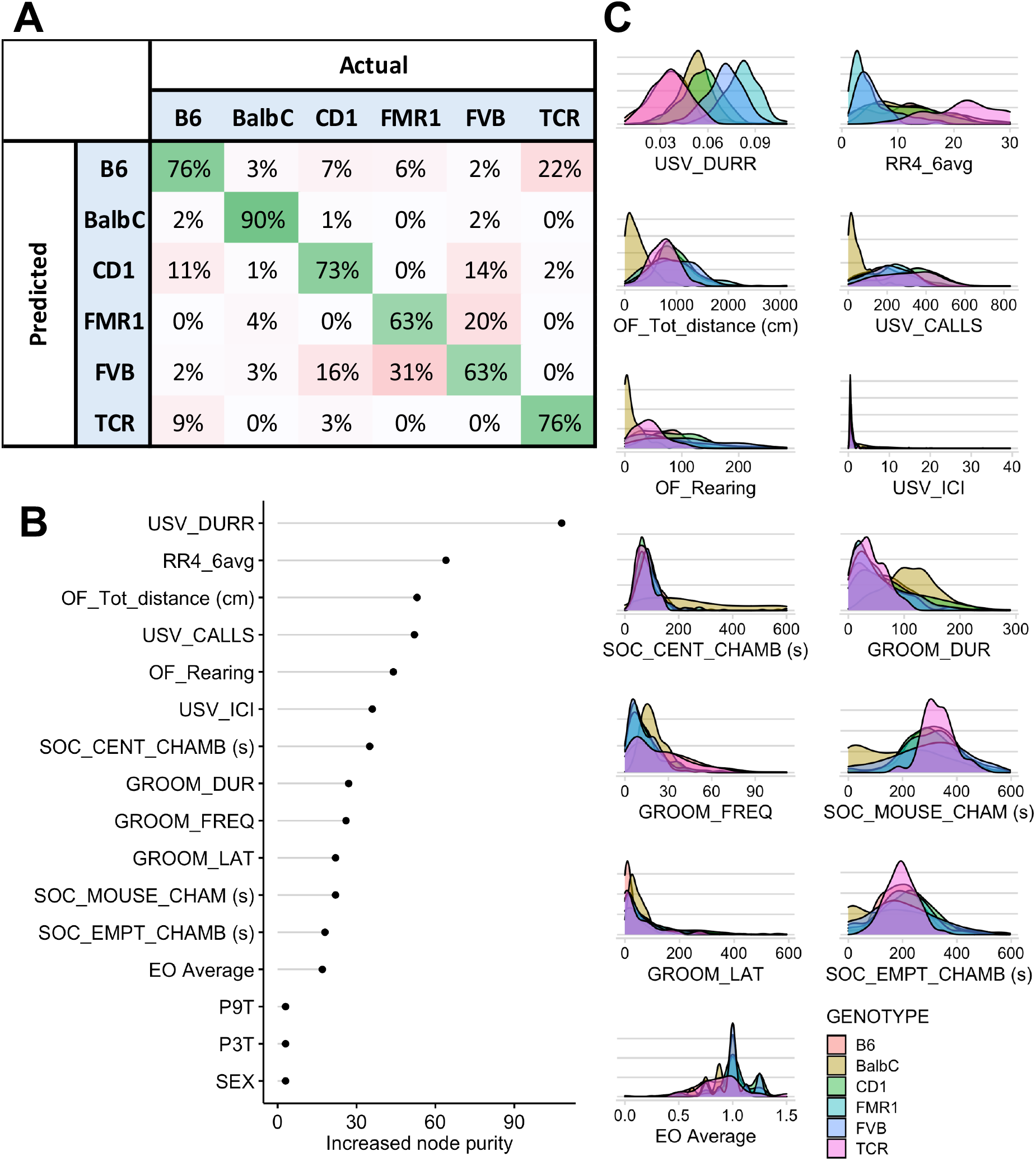
Random Forest machine learning models accurately predict mouse genotype using behavioural and neurodevelopmental outcomes. Behavioural outcomes, neurodevelopmental outcomes, sex, and treatment (P3T and P9T) were input as predictor variables. **A**) Confusion matrix represented as proportion of prediction from the total observations across 10 validation sets for each genotype. **B)** Variable importance of predictor variables for genotype as measured by increased Gini index when included in the model. **C)** Density plots of behavioural and neurodevelopmental outcomes for each mouse genotype P3T-postnatal day 3 treatment, P9T – postnatal day 9 treatment, EO – eye opening, SOC_EMP_CHAMB – time in empty chamber, SOC_CENT_CHAMB – time in the center chamber, SOC_MOUSE_CHAM – time in mouse chamber, DUR – duration, FREQ – frequency, LAT – latency, USV – ultrasonic vocalization, ICI – intercall interval, OF – open field, Tot – total, RR4 – righting reflex postnatal day 4, DURR - duration

Predictor variable importance rank and probability density function plots of behavioural and neurodevelopmental outcomes for each genotype reveals the behavioural tests and neurodevelopmental milestones that are important for genotype discrimination. USV duration was the most important predictor variable for genotype prediction across all genotypes (Fig. 5C) and had substantial area of density function separation between most genotypes (Fig. 5B). Specifically, B6 background, FVB background and Balb/C or CD1 background demonstrate visual separation of USV duration density functions (Fig. 5B). Other important predictor variables include: average righting reflex, OF total distance, USV calls, and OF rearing (Fig. 5C). Visual separation of density functions for at least one genotype exists for most of these predictor variables (Fig. 5B).

Sex and treatment were also included as predictor variables for genotype prediction in our Random Forest models, however these variables were not important predictor variables for classification of mouse genotype (Fig. 5C). This supports our ANOVA and posthoc results, which did not demonstrate any independent treatment or sex effect, and minimal treatment interactions. In our dataset, sex and LPS and/or MS treatment have limited influence on early life behavioural and neurodevelopmental trajectories. To further validate the absence of sex and treatment differences in our dataset, random forest models were trained to predict sex or treatment from behavioural and neurodevelopmental outcomes, treatment or sex, and genotype data. Accuracy for sex prediction was approximately 55%, akin to random assignment of two labels (ME = 0.45±0.04, ARI = 0.01±0.02) (Supplementary Fig. S5). Similarly, accuracy for treatment prediction was 26%, akin to random assignment of four labels (ME = 0.74±0.02, ARI = 0.001±0.014) (Supplementary Fig. S6).

### Random Forest Genotype Prediction from Relative Neuroanatomical Volumes

Mouse neuroanatomical volumes were highly associated with genotype in our study. PCA visualization of relative brain volumes from 69 regions in wildtype inbred mice leads to clustering by genotype (Fig. 6A). These visually observed clusters were validated using unsupervised hierarchical clustering (Fig. 6B). Indeed, unsupervised hierarchical clustering defines 3 highly stable clusters each dominated by a single genotype (Figure 6C). Balb/C mice have a distinct neuroanatomical phenotype that forms a pure Balb/C unsupervised cluster (Fig. 6C). Furthermore, Balb/C mice do not contaminate the B6 nor FVB neuroanatomical volume clusters. There is some overlap between the B6 and FVB clusters, with some B6 genotype mice being misclassified as FVB mice based on their relative neuroanatomical volumes. These results demonstrates that the fingerprint of mouse neuroanatomical volumes across is highly distinct for inbred mice, but more similar between B6 and FVB mice than Balb/C mice. High genotype purity of clusters indicate suggest that relative neuroanatomical volume is highly influenced by genotype.

**Figure 6:**
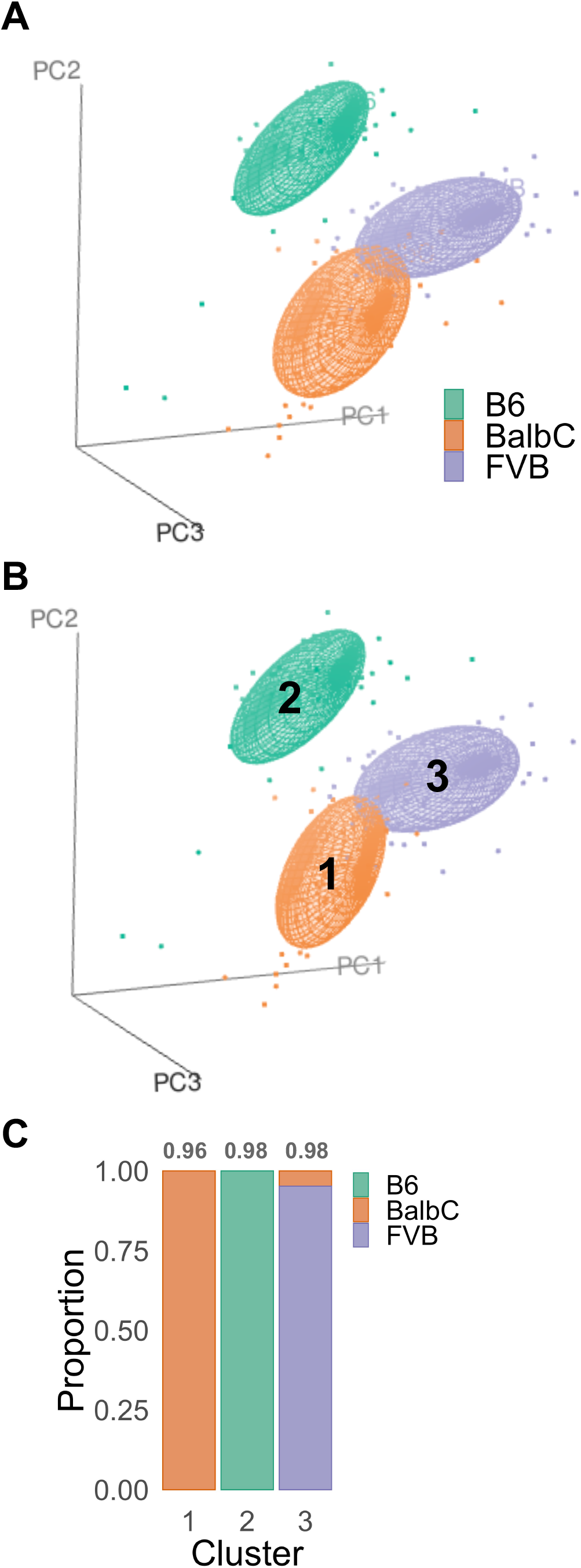
Principal component analysis of MRI relative volumes in inbred mice. Brain regions included in analysis are available in Supplementary File 3 and Supplementary Figure 7. **A)** PCA, observations coloured by Genotype. **B)** PCA points coloured by 3 clusters generated by unsupervised hierarchical clustering. Orange = Cluster 1, Green = Cluster 2, Purple = Cluster 3. **C)** Bar graph describing the proportion of observations belonging to each Genotype in Clusters I, 2, and 3.

The relationship between genotype and relative brain region volume is further supported by high accuracy of machine learning models predicting genotype from neuroanatomical volumes. Random Forest models trained using relative brain volume were 98% accurate at predicting all genotypes (ME = 0.02 ± 0.01, ARI = 0.96 ± 0.03). Misclassification was highest in FVB mice (ME = 0.05), which were most frequently misclassified as outbred CD1 mice (Fig. 7A). Inbred mice B6 and FVB and their knockout strains had very low misclassification error rates between ranging 1 – 2% (Fig. 6A), unlike the relatively higher 9 – 31% misclassification error when predicting wildtype strains from behavioural outcomes (Figure 5A). The top 20 most important brain regions for genotype discrimination were visualized as density plots (Fig. 7B, 7C). The most important brain region for genotype discrimination was Olfactory Areas (Other) – of which *TCRβ-/-δ-/-* and B6 mice have distinct distributions from the other genotypes. The caudoputamen is the next most important brain region due to high relative caudoputamen volume in Balb/C mice compared to other genotypes (Figure 7C). Overall, high genotype prediction accuracy of Random Forest models trained using relative brain volume indicates that genotype is an important factor related to individual brain region volumes.

**Figure 7:**
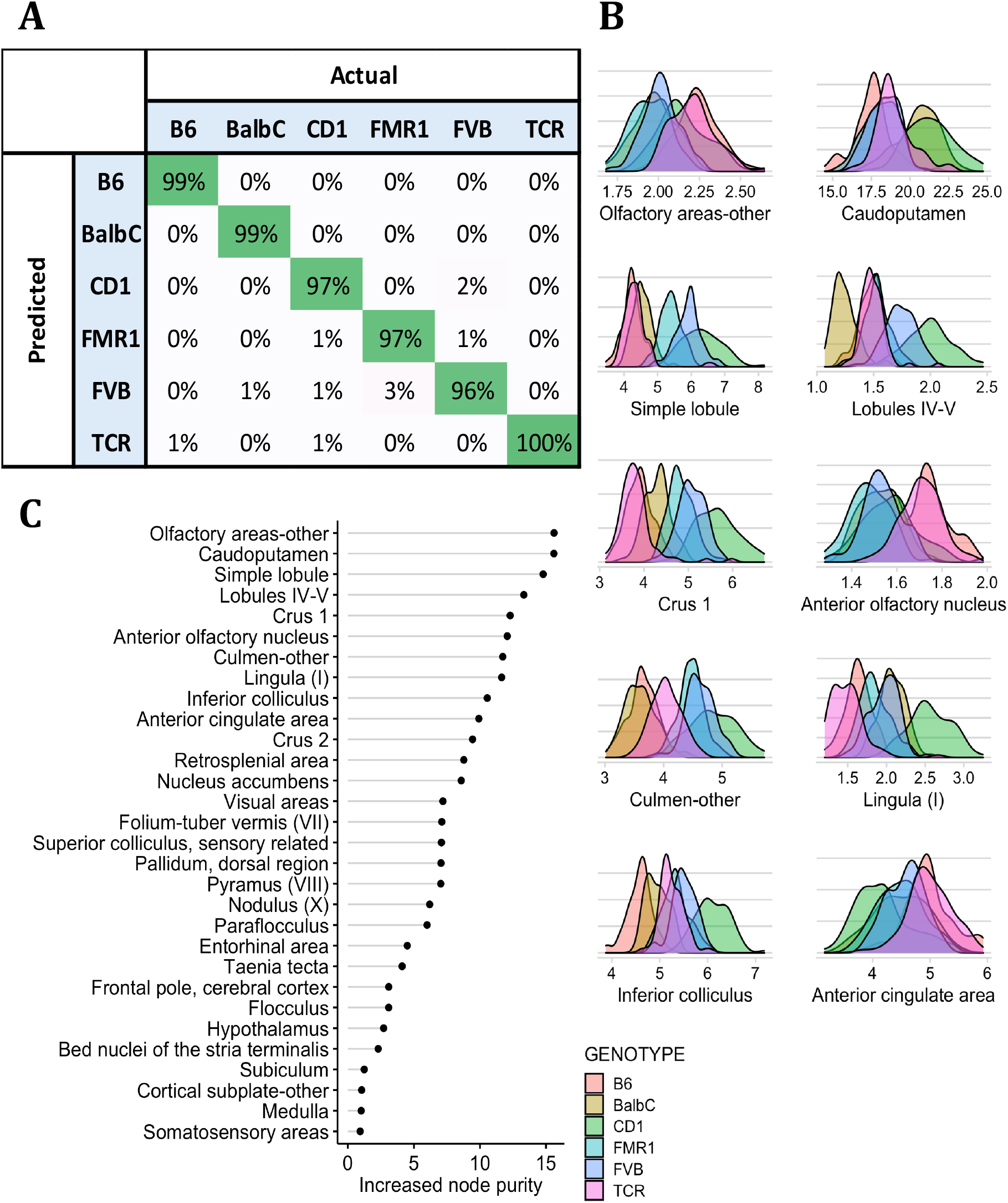
Random Forest machine learning models accurately predict mouse genotype from relative neuroanatomical volumes. The brain regions used as predictor variables are listed in Supplementary File 3. **A**) Confusion matrix represented as proportion of genotype predictions from the total number of each genotype across 10 validation sets. **B)** Variable importance of predictor variables for genotype as measured by increased Gini index when included in the model. The top 20 ranked, random 5, and lowest 5 ranked predictor variables were plotted. **C)** Density plots of brain regions in the top 20 of predictor variable importance; grouped by mouse genotype.

Random forest models were also trained to predict sex and treatment from relative neuroanatomical volumes. Random forest models predicting sex had poor prediction accuracy, only marginally better than random assignment (ME = 0.33 ± 0.04, ARI = 0.12 ± 0.06) (Supplementary Fig. 8). The Bed Nuclei of the stria terminalis was the most important predictor variable for sex. Random forest models were unable to predict treatment from relative neuroanatomical volume; prediction was akin to random assignment (ME = 0.76 ± 0.04, ARI = 0.02 ± 0.03) (Supplementary Fig. S9).

### Hierarchical Clustering of behavioural data

Hierarchical clustering was used to investigate relationships between behavioural and neurodevelopmental metrics. Clustering was performed using behavioural and milestone outcomes from mice spanning all genotypes, treatments and both sexes. Two stable clusters of neurodevelopmental milestones and behaviours emerged, indicating correlated networks of these behaviours and milestones (Fig. 8). Development metric for open field total distance was removed because of the high correlation and outcome redundancy with open field rearing. However, when open field was included in the analysis, the composition and stability the two clusters generated via hierarchical clusters were not different (Supplementary Table S1, S2). Similar behavioural networks with two highly stable clusters were identified when using inbred mice only (Supplementary Table S1, S2). The k number of clusters to generate was determined using a Silhouette plot and the elbow method interpreted form a Within-Sum-Squares (k = 2) (Supplementary Fig. S10). The dendogram in Figure 8 demonstrates the two stable clusters of behaviours and neurodevelopmental milestones. In one cluster the metrics related to grooming, righting reflex, sociability center time (movement) and intercall interval of USVs are linked. The other cluster links open field activity, eye opening, USV number and duration, and sociability. These associated networks of behaviours may indicate the neurocircuitry programming behaviours in each cluster are developmentally linked, and that mice may follow a neurodevelopmental trajectory that can be broadly characterized using two behavioural networks.

**Figure 8:**
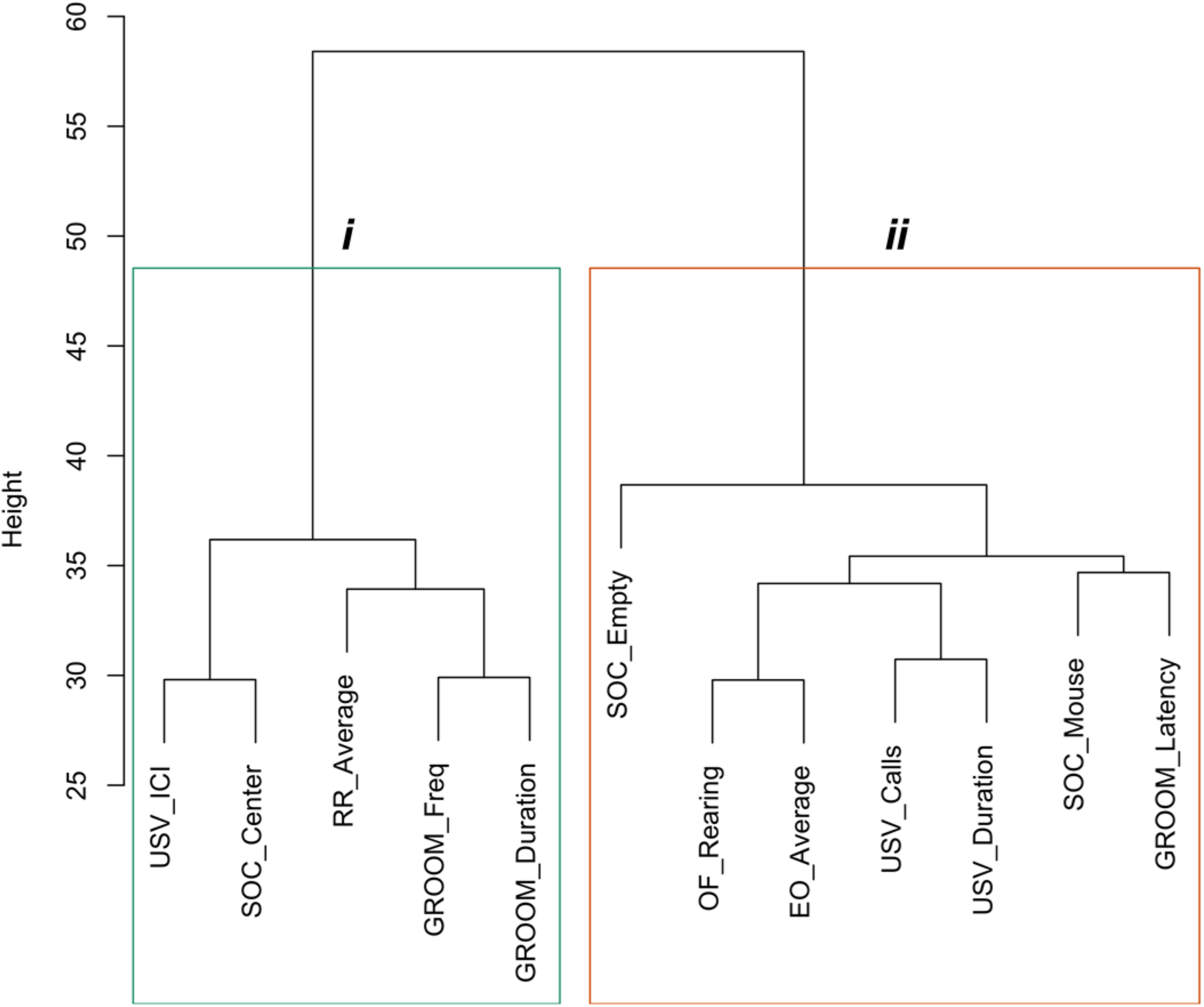
Neurodevelopment and behavioral metrics from 871 clustered by Euclidian distances. Ward.D2 hierarchical clustering method was used to generate 2 clusters. Clusters were bootstrapped to validate cluster stability (Jaccard Mean = 0.972 (Cluster i), 0.986 (Cluster ii)).

## Discussion

We have developed a 28-day pipeline that tracks the neurodevelopmental and behavioural trajectory of mice. Using this pipeline, we assessed the influence of genetics, early-life challenge, and sex on the behavioural and neurodevelopmental trajectory for over 1060 mice. To our knowledge, our study is the first big data mouse neurodevelopment project using clinically relevant developmental milestones and behavioural tests. While other behavioural neuroscience studies have used large sample sizes, the sample sizes are often diluted across experimental groups, preventing the use of statistical methods requiring large sample sizes for accurate analyses, such as hierarchical clustering and machine learning class predictions.

There are logistical challenges to curating a large-scale behavioural mouse database. A Big Data Mouse pipeline is vulnerable to issues of reproducibility and test validity. Each mouse in the data collection pipeline must undergo the same experimental tests at specific time points, as behavioural test results in mice are impacted by developmental age and the order of behavioural tests. Furthermore, errors arise from collecting data over an extended period, therefore it is essential there are robust quality control protocols ensuring data collected from daily assessments and tests are accurately transferred to the database, and automated behavioural test equipment is used whenever possible.

Our findings demonstrate the significant influence of background genetics on behaviour and neurodevelopmental trajectory. The main effects on neurodevelopmental milestones and behaviour tests identified in the between-subjects’ effects were predominantly genotype differences. Posthoc analysis further revealed that differences in behaviour often existed between mouse strains, and less frequently between KO mice and their wild type counterparts. Inbred mouse strain differences in specific behaviours and developmental milestones culminate into to a neurodevelopmental phenotype unique to each strain, which can be identified via differential clustering on PCA plots. Hierarchical Clustering validates visually observed inbred strains clusters, with each unsupervised cluster was dominated by one inbred strain. Random Forest models trained using neurodevelopmental and behavioural data accurately predict genotype, which further validates distinct neurodevelopmental and behavioural phenotypes associated with each genotype. Machine learning model misclassifications typically occurred between knockout mice and their background strains, or inbred mice and outbred CD1 mice, demonstrating the phenotypic overlap between these genotypes. It is likely genetic diversity between individual outbred CD1 mice contributes to the heterogeneous CD1 neurodevelopmental phenotype, which varied between the FVB phenotype and a subset of the B6 neurodevelopmental phenotype in our study. While the relationship between behaviour and genetics in adult mice has already been robustly established, our study demonstrates how genetic background, and consequently behavioural outcomes, define the range of overall early-life behavioural and neurodevelopmental trajectory. Indeed, each mouse genotype could be broadly classified by a unique behavioural and neurodevelopmental phenotype.

Knockout of key neurodevelopment-associated genes produce diverse behavioural and neurodevelopmental phenotypes dependent on the background strain (Born et al., 2017; Huang et al., 2013; Lai et al., 2016; Lai et al., 2014; Pietropaolo et al., 2011; Samaco et al., 2013; Spencer et al., 2011). Indeed, hierarchical clustering predominantly grouped knockout mice with their background strain in the PCA analysis. Random Forest models trained to predict genotype further validated this relationship, as knockout mice were primarily misclassified as their own background strain. Although knockout mice have differ from their background strain in select behavioural tests and neurodevelopmental milestones, high-dimensional analyses using machine learning demonstrate that the phenotype of knockout mice broadly intersects with mice in their background strain. Our study validates the relevance of the background strain on behavioural phenotype in knockout mice models, as the background strain is the primary determinant of overall neurodevelopmental phenotype.

We also demonstrate robust neuroanatomical phenotypes delineated by mouse strain and knockout genotype. Random Forest models trained using relative brain region volume predict mouse strain and knockout genotype with 98% accuracy, suggesting that neuroanatomical volumes were largely driven by genotype. Additionally, both knockout strains can be differentiated from their background strain, indicating the FMR1 gene (*FMR1-KO* model) and T cells (*TCRβ-/-δ-/-* model) influence neuroanatomical volumes in adolescence. Significant differences between mouse strains’ neuroanatomical volume and morphological variance have been previously documented in several studies (Chen et al., 2006; Lin et al., 2015; Scholz et al., 2016). Additionally, quantitative trait loci that influence neuroanatomical volume have been identified, indicating the role of genetics (Rosen & Williams, 2001; Zygourakis & Rosen, 2003). We build upon existing literature by using machine learning models to demonstrate that neuroanatomical volume differences between mouse genotypes are profound enough for near-perfect supervised and unsupervised classification.

Unlike early-life behavioural phenotype in knockout mice, which were most frequently misclassified as their background strain, there is high purity when classifying early-life neuroanatomical volumes of knockout mice (*FMR1-KO* and *TCRβ-/-δ-/-* model) and their background strains (FVB and B6). These results suggest a disconnect between neuroanatomical differences and behavioural differences that may be driven by background genetics. Broadly, neuroanatomical diversity may be primarily dictated by genotype whereas behavioural phenotypic diversity is likely to be more susceptible to gene-environment interactions. The observed genotypic stability of neuroanatomical phenotypes may have translational applications for diagnosing subtypes of neurodevelopmental disorders with predominantly genetic causes. Indeed, significant neuroanatomical volume differences have been observed in several neurodevelopment disorders including attention deficit hyperactivity disorder and autism spectrum disorder (Boedhoe et al., 2020; Hazlett et al., 2012; Hoogman et al., 2017; Lin et al., 2015).

Finally, we provide evidence for two highly stable networks of connected early-life behaviours and neurodevelopmental milestones. The existence of these networks implies that early developmental milestones may accurately predict behavioural phenotypes later in life related to behavioural activation and inhibition. We identified two networks of behavioural outcomes positively correlated to neurodevelopmental milestones using hierarchical clustering of behavioural parameter principal components. Cluster 1 can be summarized as behavioural inhibition. Righting reflex is the developmental milestones associated with Cluster 1, and behaviours include: grooming frequency and duration, 3-chambered sociability center time, and USV inter-call interval. The behaviours in the Cluster 1 are repetitive or represent low engagement for early life and adolescent social parameters. The cluster 1 milestone – righting reflex – also represents behavioural inhibition, as higher average righting reflex score indicates delayed motor development. Late onset of motor development in mice may be predictive of an inhibited behavioural phenotype later in life. Cluster 2 can be summarized as behavioural activation and is associated with the eye-opening milestone. The behaviours of Cluster 2 include: USV Calls and duration, 3-Chambered Sociability Mouse or Empty chamber time, Open-field parameter, and latency to groom. The behavioural readouts are proxies for high locomotor and social activity throughout early and adolescent life. Mice who achieve the eye-opening score earlier in development may be more likely to have an activated behavioural phenotype in adolescence. Indeed, early life behavioural milestones achievement have clinical associations with behavioural trajectory, in particular, for early detection of neurodevelopmental disorders in clinical populations (Gurevitz et al., 2014; Havdahl et al., 2021; Johnson et al., 2015; Provost et al., 2007). The discovery of milestone-associated behavioural networks in our study indicates the external validity of our behavioural pipeline for modelling neurodevelopmental trajectories.

In summary, we developed a behavioural and neurodevelopmental pipeline to demonstrate the influence of genetic background on neurodevelopmental trajectory and neuroanatomical phenotype, as well as the relationship between neurodevelopmental milestones and behavioural outcomes. Furthermore, our pipeline contextualizes the effect of knockouts targeting neurodevelopmental on behavioural outcomes that differ by strain. By applying hierarchical clustering to neurodevelopmental data of knockout mice and their background strain, the broad similarity of their neurodevelopmental trajectory can be assessed. Detection of robust genotype effects for behaviours and neuroanatomy, as well as early neurodevelopmental milestones being predictive of adolescent behavioural trajectory, indicates the external validity of our behavioural pipeline. Ultimately, this standardized neurodevelopmental trajectory pipeline could be used to model how genotype, environmental insults, or drug interventions influence clinically relevant neurodevelopmental and behavioural outcomes throughout adolescent development.

