## Supplemental Figures for "Host genetics maps to behaviour and brain structure in mice"

Supplemental Figures and Tables

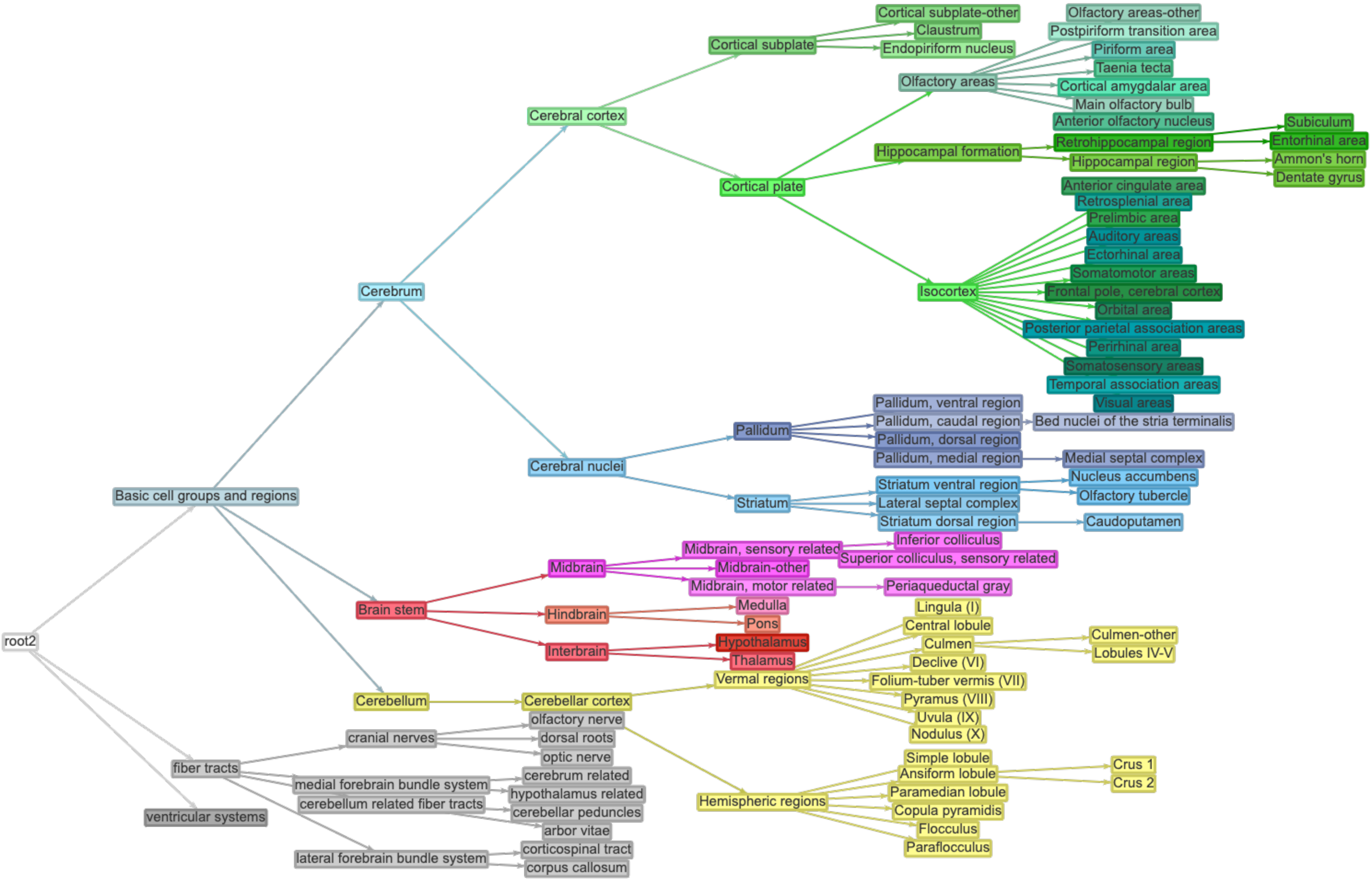


**Supplementary Figure S1**: Pruned hierarchical anatomy of brain regions. Leaves of the hierarchical tree (also listed in Supplementary File 1) were used for neuroanatomical PCA or random forest analyses.

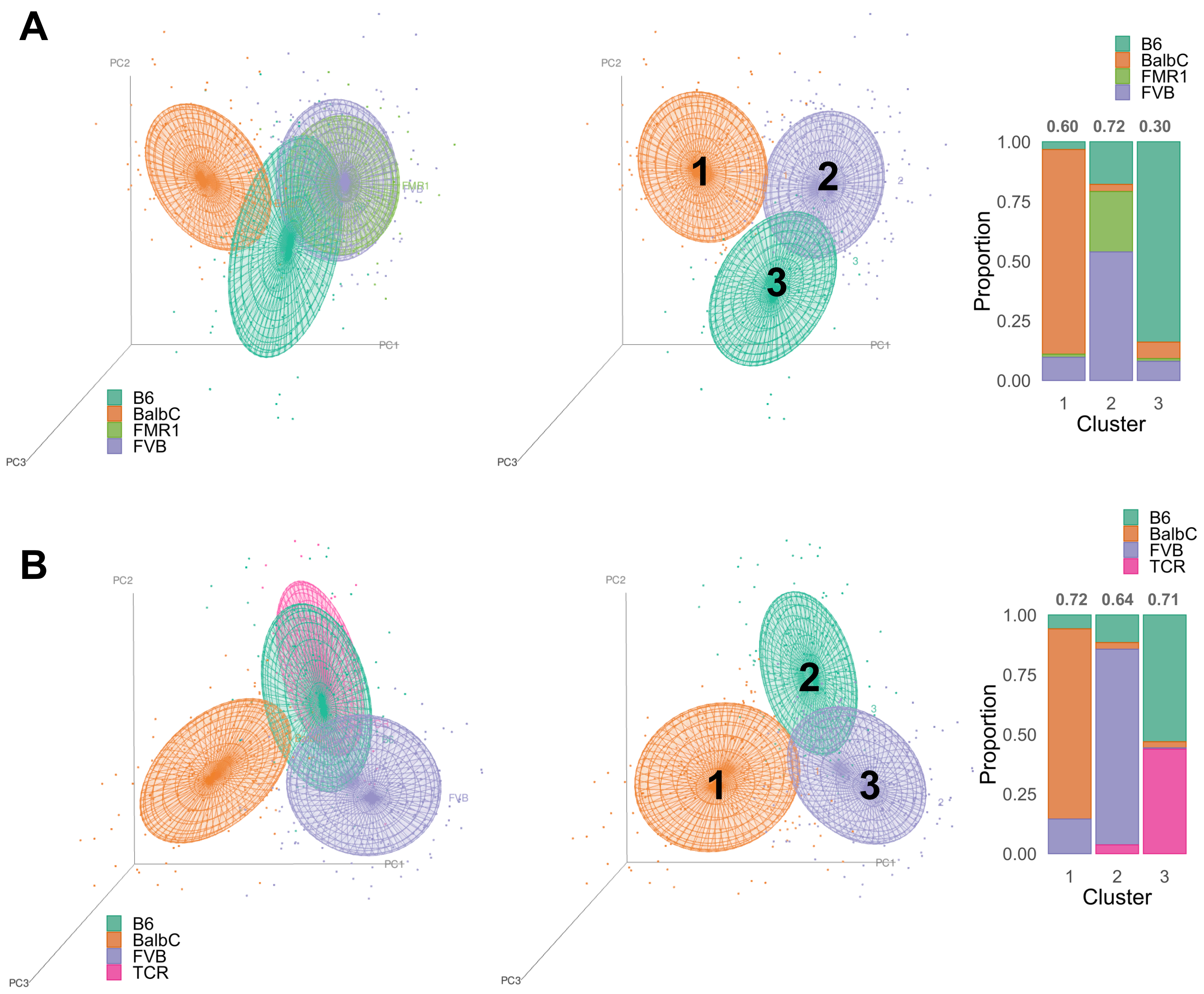


**Supplementary Figure S2**: Neurodevelopment and behavioral PCA for inbred and knockout mice. **A)** PCA plot of inbred mice and *FMR1-KO* mice clustered by genotype **(left)** or unsupervised hierarchical clustering **(middle)**. Bar chart **(right)** of the behavioural clusters composition by genotype. Cluster stability – as measured by Jaccard Index – is given above the bar for each cluster. **B)** PCA plot of inbred mice and *TCRβ-/-δ-/-* mice clustered by genotype **(left)** or unsupervised hierarchical clustering **(middle)**. Bar chart **(right)** of the behavioural clusters composition by genotype. Cluster stability – as measured by Jaccard Index – is given above the bar for each cluster.

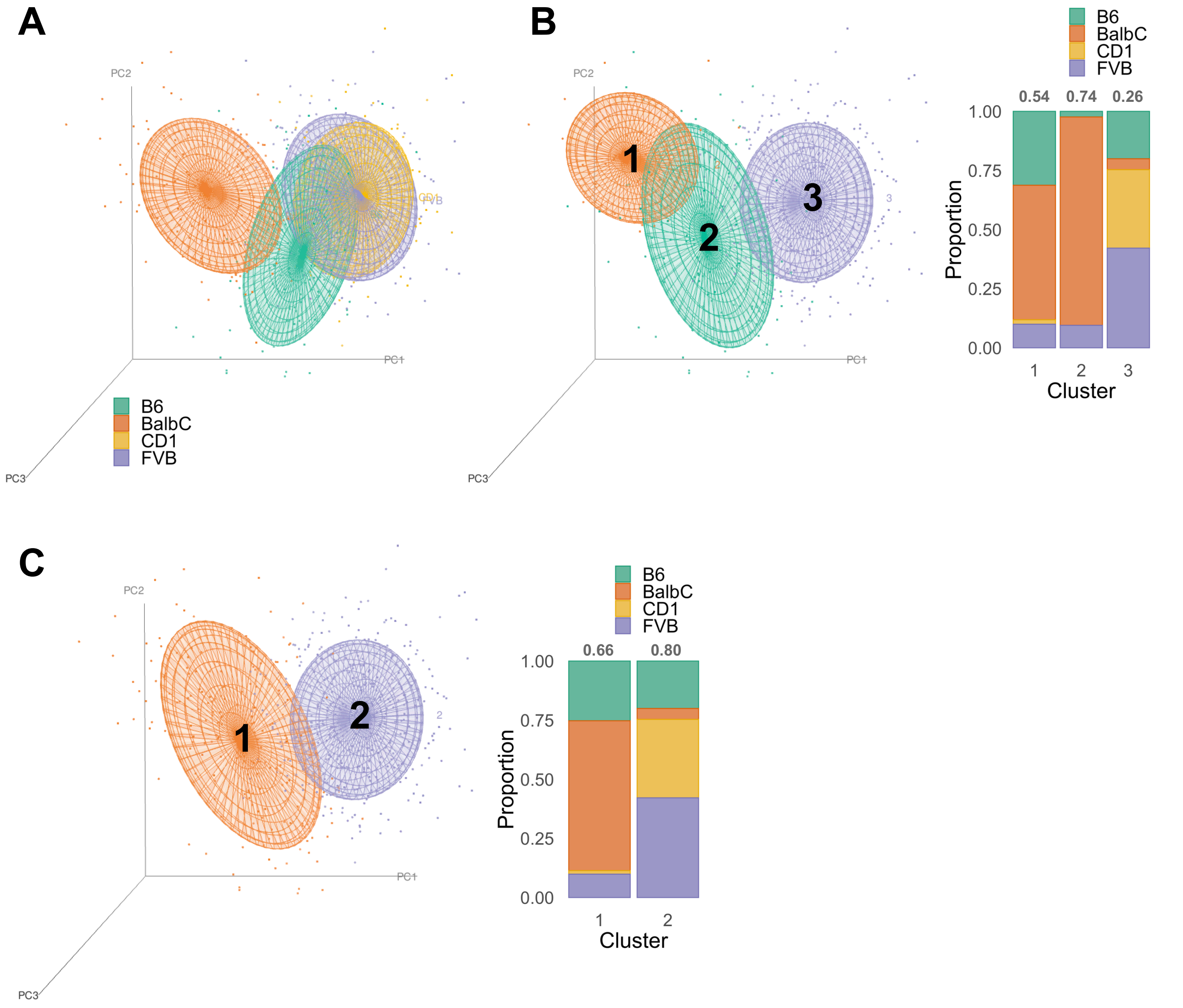


**Supplementary Figure S3:** Neurodevelopmental and behavioral PCA in inbred and outbred (CD1) mice. **A)** PCA plot of inbred mice and CD1 mice clustered by genotype. **B)** PCA plot of inbred and CD1 mice clustered by hierarchical clustering (k = 3 clusters). Bar chart of the behavioural clusters composition by genotype. Cluster stability – as measured by Jaccard Index – is given above the bar for each cluster. **C)** PCA plot of inbred and CD1 mice clustered by hierarchical clustering (k = 2 clusters). Bar chart of the behavioural clusters composition by genotype. Cluster stability – as measured by Jaccard Index – is given above the bar for each cluster.

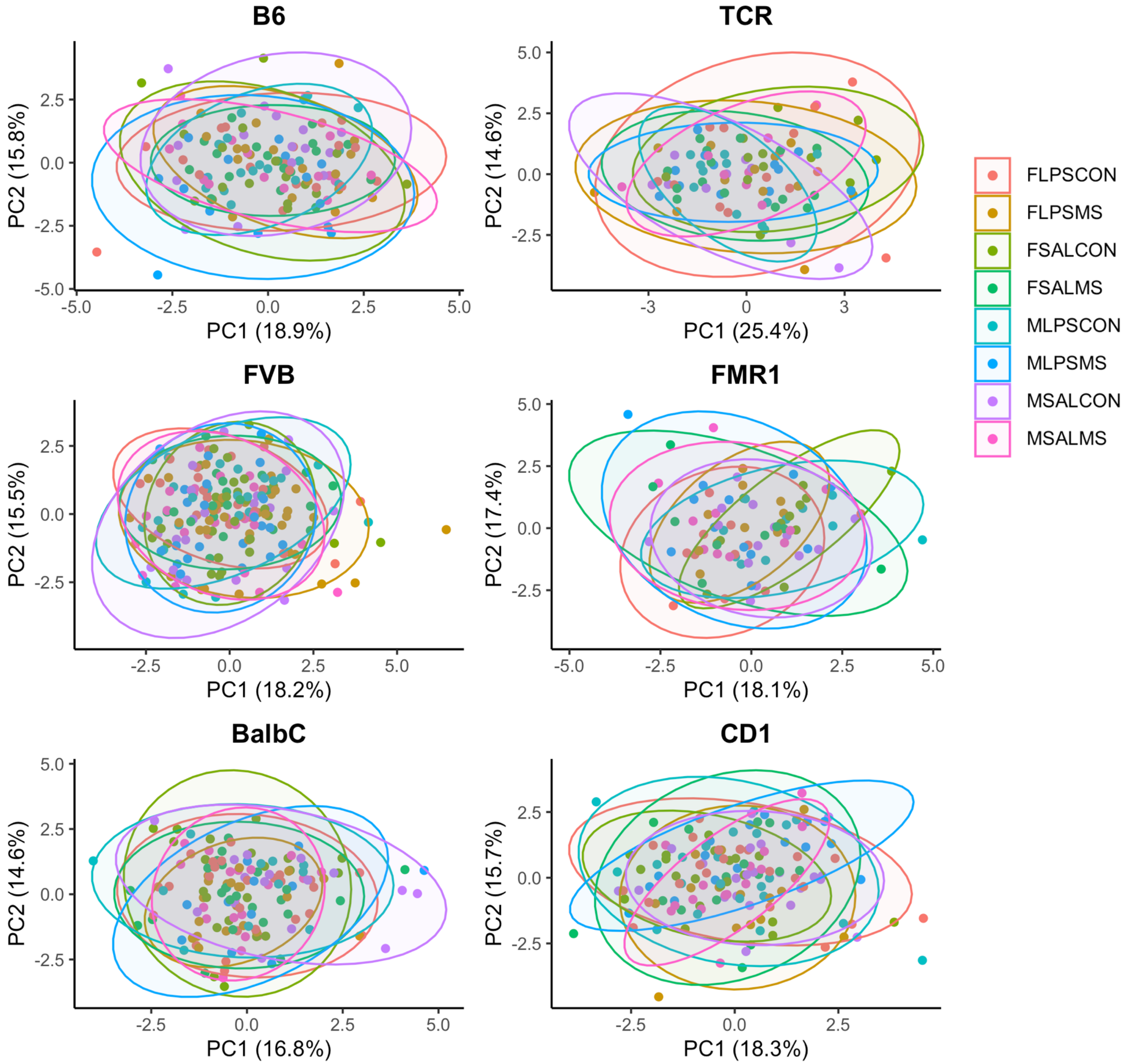


**Supplementary Figure S4:** 2D principal component analysis suggests limited differences in neurodevelopmental trajectory when mice are grouped by combined treatment and sex within each genotype. F = Female or M = Male. SAL = saline control (P3) or LPS = LPS treated mice (P3). MS = maternal separation (P3) or CON = non-maternally separated (P9).

**A**

| | | Actual | |
|---|---|---|---|
| | | F | M |
| Predicted | F | 58% | 48% |
| | M | 42% | 52% |

**B**

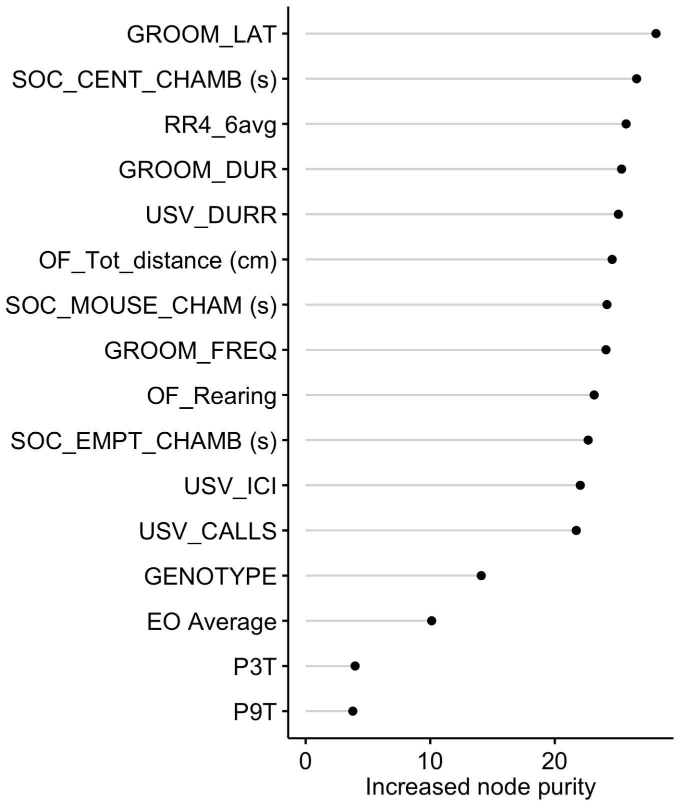


**C**

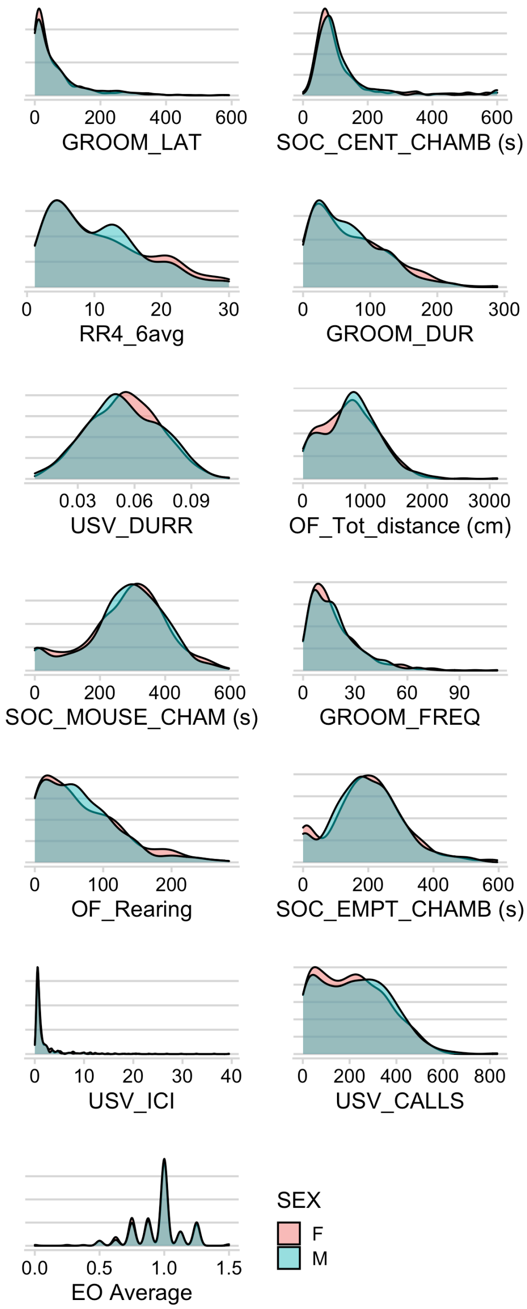


**Supplementary Figure S5:** Random Forest model predicting sex using neurodevelopmental and behavioural outcomes, treatment, and genotype. **A)** Classification matrix represented as proportion of predicted sex from total of the female or male samples input across 10 validation sets. **B)** Variable importance of random forest predictor variables for sex as measured by increased Gini index when included in the model. **C)** Density plots demonstrating behavior and neurodevelopmental milestone distributions for each sex.

**A**

| | | Actual | | | |
|---|---|---|---|---|---|
| | | LPS_CON | LPS_MS | SAL_CON | SAL_MS |
| Predicted | LPS_CON | 25% | 23% | 25% | 26% |
| | LPS_MS | 21% | 29% | 26% | 26% |
| | SAL_CON | 32% | 20% | 23% | 22% |
| | SAL_MS | 22% | 29% | 26% | 26% |

**B**

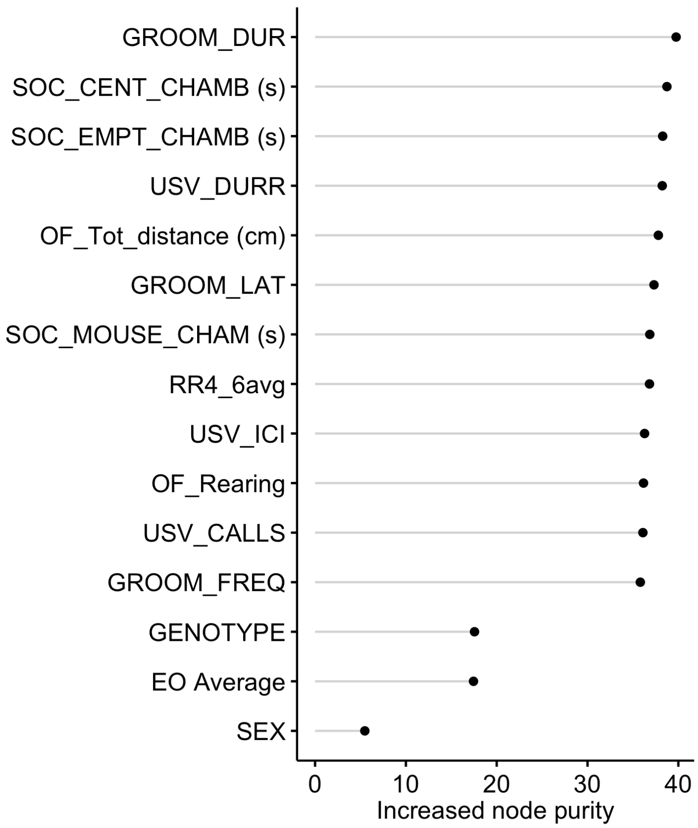


**C**

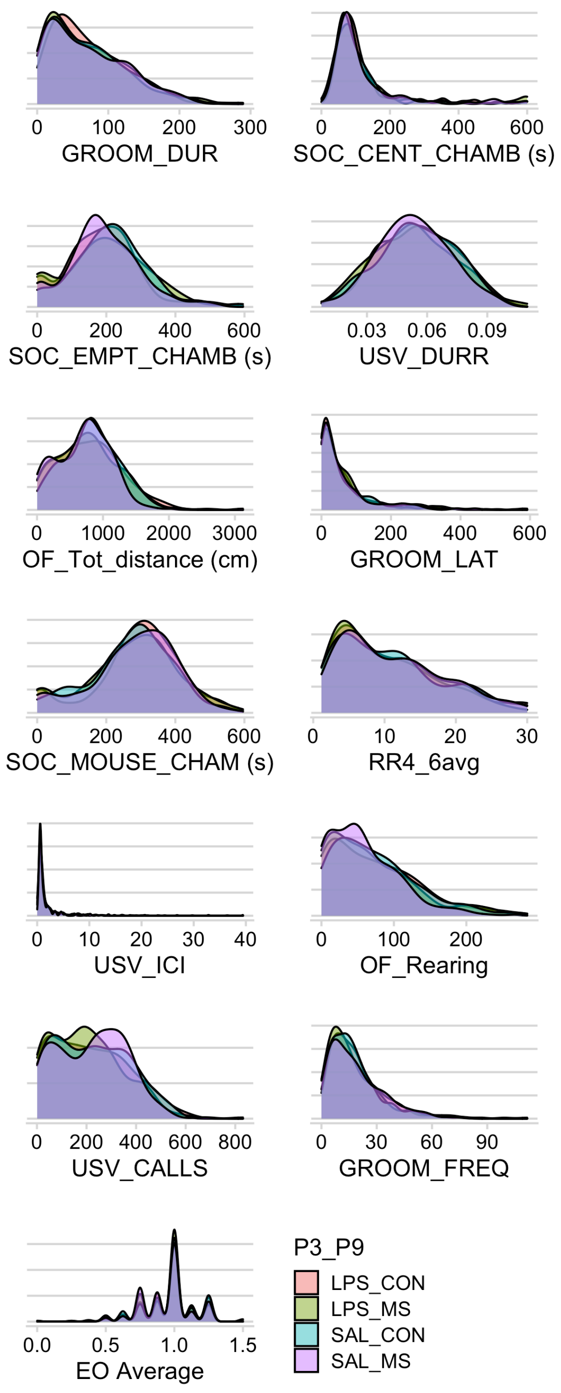


**Supplementary Figure S6:** Random Forest model predicting combined treatment using neurodevelopmental and behavioural outcomes, treatment, and genotype. **A)** Classification matrix represented as proportion of predicted treatment from total of the female or male samples input across 10 validation sets. **B)** Variable importance of random forest predictor variables for treatment ranked by increased Gini index when included in the model. **C)** Density plots demonstrating behavior and neurodevelopmental milestone distributions for each treatment.

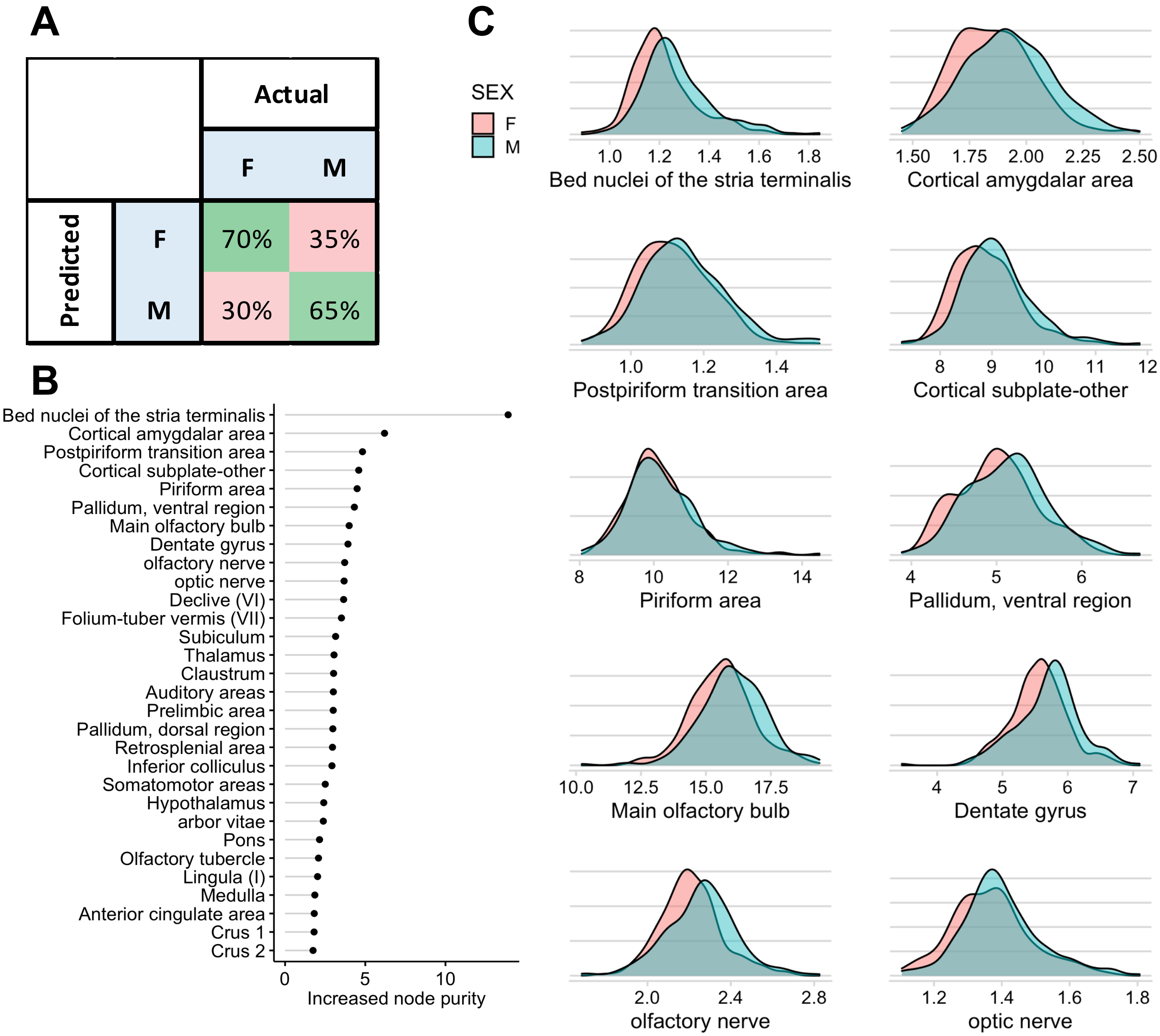


**Supplementary Figure S8:** Random Forest machine learning models had low accuracy in predicting sex from relative neuroanatomical volumes. Brain regions used as predictor variables are listed in Supplementary File 3. **A)** Confusion matrix represents proportion of predicted sex from total of male or female observations to predict across 10 validation sets. **B)** Variable importance of predictor variables for sex as measured by increased Gini index when included in the model. The top 20 ranked, random 5, and lowest 5 ranked predictor variables were plotted. **C)** Density plots of brain regions in the top 20 of predictor variable importance; grouped by sex.

**A**

| | | Actual | | | |
|---|---|---|---|---|---|
| | | LPS_CON | LPS_MS | SAL_CON | SAL_MS |
| Predicted | LPS_CON | 0.28 | 0.15 | 0.36 | 0.21 |
| | LPS_MS | 0.16 | 0.25 | 0.21 | 0.35 |
| | SAL_CON | 0.31 | 0.22 | 0.22 | 0.23 |
| | SAL_MS | 0.26 | 0.38 | 0.21 | 0.22 |

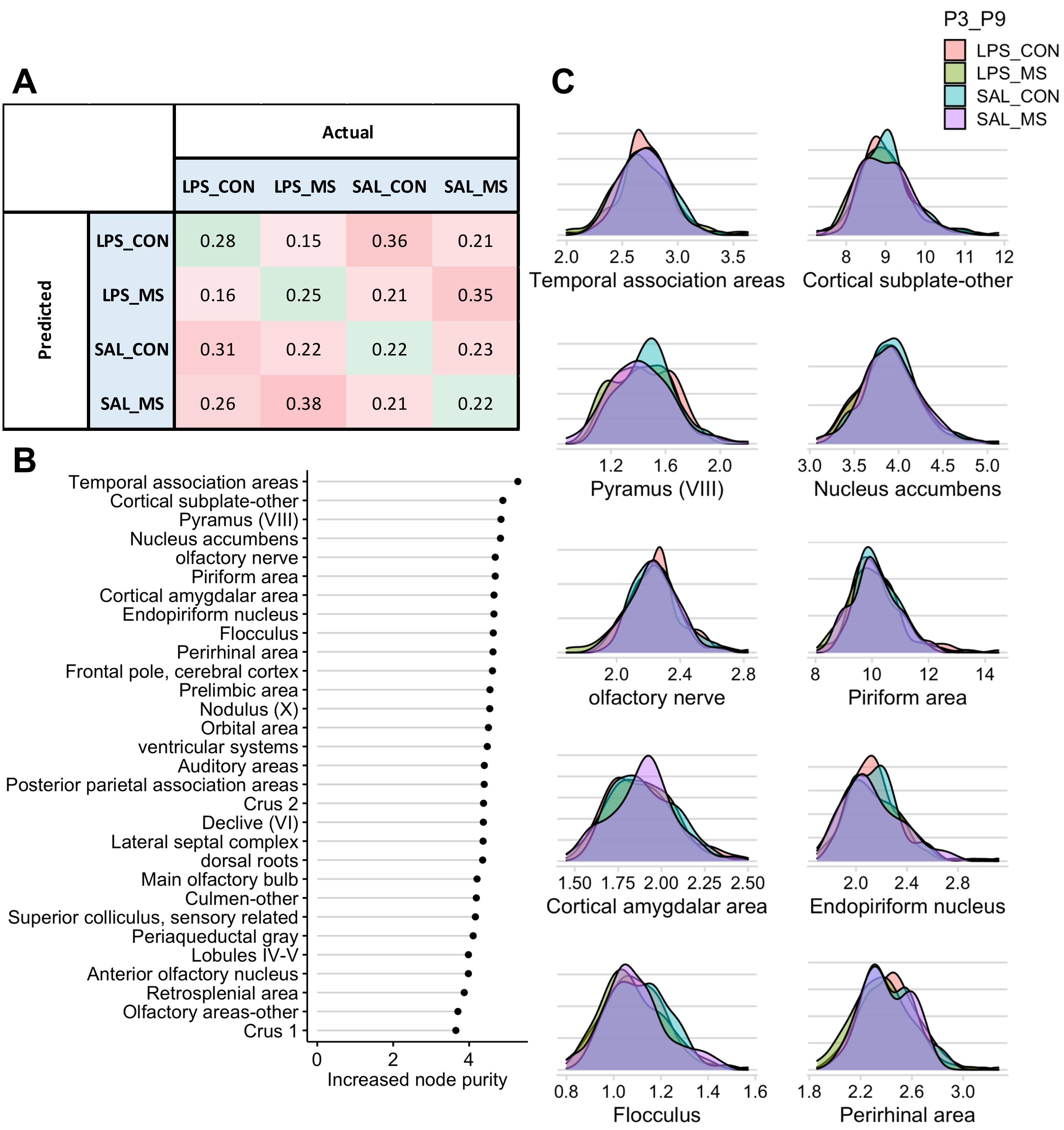


**Supplementary Figure S9:** Random Forest machine learning models were unable to predict early-life stress treatment from relative neuroanatomical volumes. Brain regions used as predictor variables are listed in Supplementary File 3. **A**) Confusion matrix represents proportion of predicted treatment from total of the treatment class input across 10 validation sets. **B)** Variable importance of predictor variables for treatment as measured by increased Gini index when included in the model. The top 20 ranked, random 5, and lowest 5 ranked predictor variables were plotted. **C)** Density plots of brain regions in the top 20 of predictor variable importance; grouped by early-life stress treatment.

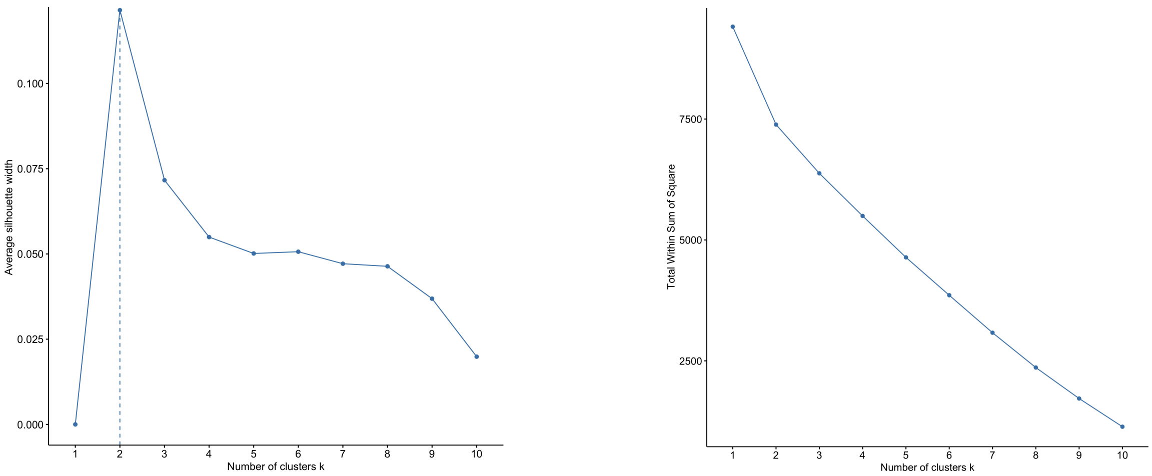


**Supplementary Figure S10:** Silhouette plot and Elbow plot for determining optimal number of behavioural and neurodevelopmental clusters. 2 clusters (k) were selected for hierarchical clustering.

| **Cluster** | **1** | **2** |
|---|---|---|
| **All Genotypes** | 0.906 | 0.950 |
| **Wildtype** | 0.965 | 0.982 |
| **AllGenotypes_OFdist** | 0.956 | 0.967 |
| **AllGenotypes_OFrear** | 0.937 | 0.957 |
| **Wildtype_OFdist** | 0.980 | 0.987 |
| **Wildtype_OFrear** | 0.972 | 0.986 |

**Table S1:** Stability of behavioural networks was validated using cluster stability analysis. Cluster stability is reported as the mean Jaccard Index from bootstrapped samples. Network iterations are labelled using the following terms: **AllGenotypes** = BalbC, B6, FVB, CD1, FMR1-KO, TCRβ-/-δ-/- strains used to generate behavioural cluster. **Wildtype** = BalbC, B6, FVB strains used to generate behavioural clusters. **OFdist** = Only open field total distance metric included in analysis, i.e. open field rearing metric discarded before generating behavioural cluster. **OFrear** = Only open field rearing metric included included in analysis, i.e. open field total distance metric discarded before generating behavioural cluster.

| Behaviour | All Genotypes | Wildtype | AllGenotypes_OFdist | AllGenotypes_OFrear |
|---|---|---|---|---|
| Average Righting Reflex | 1 | 1 | 1 | 1 |
| USV Calls | 2 | 2 | 2 | 2 |
| USV ICI | 1 | 1 | 1 | 1 |
| USV Duration | 2 | 2 | 2 | 2 |
| OF Rearing | 2 | 2 | NA | 2 |
| OF Total Distance | 2 | 2 | 2 | NA |
| Soc Center Chamber | 1 | 1 | 1 | 1 |
| Soc Empty Chamber | 2 | 2 | 2 | 2 |
| Soc Mouse Chamber | 2 | 2 | 2 | 2 |
| Groom Frequency | 1 | 1 | 1 | 1 |
| Groom Duration | 1 | 1 | 1 | 1 |
| Groom Latency | 2 | 2 | 2 | 2 |
| Eye Opening Score | 2 | 2 | 2 | 2 |

**upplementary Table S2:** Cluster assignment of each behaviour or neurodevelopmental
ilestone in each iterations of the behavioural clustering analysis. Network iterations are labelled
sing the following terms: **AllGenotypes** = BalbC, B6, FVB, CD1, FMR1-KO, TCRβ-/-δ-/- strains
sed to generate behavioural cluster. **Wildtype** = BalbC, B6, FVB strains used to generate
ehavioural clusters. **OFdist** = Only open field total distance metric included in analysis, i.e. open
eld rearing metric discarded before generating behavioural cluster. **OFrear** = Only open field
aring metric included included in analysis, i.e. open field total distance metric discarded before
enerating behavioural cluster.
